# Global gradients in species richness of marine plankton functional groups

**DOI:** 10.1101/2023.07.03.547473

**Authors:** Fabio Benedetti, Nicolas Gruber, Meike Vogt

## Abstract

The patterns of species diversity of plankton functional groups (PFGs) remain poorly understood although they matter for marine ecosystem functioning. Here, we use an ensemble of empirical species distribution models for 845 plankton species to estimate the mean annual global species richness of three phytoplankton and eleven zooplankton functional groups as a function of objectively selected environmental predictors. The species richness of all PFGs decreases from the low to the high latitudes, but the steepness and the shape of this decrease varies significantly across PFGs. Pteropods, small copepods (Oithonids and Poecilostomatoids) and Salps show the steepest latitudinal gradients, whereas Amphipods and the three phytoplankton groups show the weakest ones. While the position of such peaks and troughs in richness is modulated by the presence of upwelling systems, boundary currents and oxygen minimum zones, the gradients of temperature, irradiance, and nutrient concentration are the first-order control on the main latitudinal richness patterns. The species richness of all PFGs increases with net primary production but decreases with particles size and the efficiency of the biological carbon pump. Our study puts forward emergent Biodiversity-Ecosystem Functioning relationships and hypotheses about their underlying drivers for future field-based and modelling research.

## 1. Introduction

Countless micro- and macroplankton drive biogeochemical cycles and ecosystem processes in the oceans (Falkowski & Oliver, 2007; Landry & Steinberg et al., 2017). Phytoplankton (i.e., the photoautotrophic microalgae and bacteria) carry out nearly half of the planetary net primary production (Field et al., 1998), thereby fueling the transfer of energy from the basis of all marine ecosystems to higher trophic levels. By doing so, phytoplankton control the concentration and distribution of nutrients (Martiny et al., 2013; Inomura et al., 2022). Phytoplankton represent a major food source for the zooplankton (i.e., the heterotrophic protists and animals) which ensure secondary production and sustain global fisheries (Beaugrand et al., 2010; Landry & Steinberg et al., 2017). Together, phytoplankton and zooplankton modulate the biogeochemical carbon pump which helps regulating Earth’s climate by sequestrating atmospheric CO_2_ (Sarmiento & Gruber, 2006). These major ecosystem functions are insured by more than 100’000 different species altogether, some of which remain unknown in spite of centuries of research (de Vargas et al., 2015; Abreu et al., 2022).

While we do not know all plankton species, we know that variations in evolutionary histories lead to major inter-group and inter-species differences in functional traits and thus different contributions to ecosystem functions (Litchman et al., 2009, 2013; Henson et al., 2019; Martini et al., 2021). Major differences in cell size (or body size for multicellular organisms), morphology, elemental composition, feeding modes, life cycles or behaviors exist across and within the extant taxonomic groups of the plankton (Beaugrand et al. 2010; Assmy et al., 2013; Henschke et al. 2016; Brun et al., 2017; McConville et al., 2017; Conley et al., 2018; Tréguer et al., 2018). For instance, diatoms take up silicic acid to build silicified cell walls that protect them from grazers and increase their sinking rates (Tréguer et al., 2018), a trait that is unique in the phytoplankton. In spite of this group-specific, unifying trait, diatom species exhibit a variety of cell size, morphologies, silica content and life strategies that affect diatom population dynamics, nutrients uptake rates and carbon export fluxes (Assmy et al, 2013; Leblanc et al., 2018; Tréguer et al., 2018). In the zooplankton, copepods alone include species of body lengths varying from 0.1 to 10 mm, with passive or active feeding techniques that modulate their grazing and mortality rates (Benedetti et al., 2016; Brun et al., 2017; van Someren Grève et al., 2017). Phytoplankton-grazing crustacean zooplankton are made up of more carbon than carnivorous and flux-feeding gelatinous clades such as jellyfish and tunicates (Kiørboe, 2013; McConville et al., 2017; Conley et al., 2018). Hence, inter- and intra-group species diversity and biotic interactions matter for marine ecosystem functioning on local to global scales (Strong et al., 2015; Guidi et al., 2016; Gonzalez et al., 2020). However, the patterns of plankton species diversity and their underlying drivers remain poorly understood, despite of more than 60 years of research (Hutchinson, 1959).

To deal with such biological complexity, functional types emerged as a concept to aggregate tens of thousands of existing plankton taxonomic units (e.g. species) into fewer manageable compartments to represent the role of plankton in mechanistic ecosystem models (Le Quéré et al., 2005; Hood et al., 2006). However, most of these mechanistic models are not designed to capture the vast diversity of species and traits that exists within functional types (Anderson, 2005; Le Quéré et al., 2005) and they are not designed to study the processes that generate plankton species diversity patterns, except for a few (Barton et al., 2010; Ward et al., 2012; Dutkiewicz et al., 2020). Field observations remain essential to document the diversity patterns of plankton functional types so we can unravel their abiotic drivers and then improve and benchmark new ecosystem models that better accommodate biological diversity. Recent empirical modelling studies showed that phyto- and zooplankton display latitudinal diversity gradients that resemble the ones of other marine ectotherms, with increasing number of species from the high to the low latitudes (Righetti et al., 2019; Ibarbalz et al., 2019; Benedetti et al., 2021), a pattern mainly driven by temperature (Tittensor et al., 2010). Other observational and model-based studies reported a much weaker role of temperature in shaping phytoplankton diversity (Rodriguez-Ramos et al., 2015; Busseni et al., 2020; Dutkiewicz et al., 2020) and emphasized the role of mixing and nutrient concentrations (Barton et al., 2010; Vallina et al., 2014; Mangolte et al., 2022). Within trophic levels, observation-based studies documented global and regional latitudinal diversity gradients for multiple taxonomic groups and/or functional types: Diatoms (Malviya et al., 2017; Endo et al., 2018; Busseni et al., 2020), Coccolithophores (O’Brien et al., 2016; Endo et al., 2018); Dinoflagellates (Chen et al., 2011; Le Bescot et al., 2016), Microzooplankton (Dolan et al., 2016), Copepods (Wood-Walker et al., 2002; Rombouts et al., 2011; Hirai et al., 2020), Euphausiids (Tittensor et al., 2010), Foraminifera (Lombard et al., 2011; Yasuhara et al., 2020; Ruffino et al., 2022), Radiolaria (Boltovskoy & Correa, 2017; Biard et al., 2017), Chaetognaths (Miyamoto et al., 2014), Pteropods (Burridge et al., 2016), and hyperiid Amphipods (Burridge et al., 2017). The broad variety of scales, sampling methodologies and numerical approaches covered by these studies make inter-group and intra-group comparisons of diversity patterns difficult and differences very hard to explain. Because some groups are inadequately sampled by traditional sampling techniques (i.e., plankton nets), their contributions to community abundance, biomass and diversity have been historically underestimated, and global estimates of species diversity and composition are still missing for many key PFGs (i.e., Dinoflagellates, Ciliates, Pteropods, Tunicates, Jellyfish, Chaetognaths or Amphipods). A major consequence of these historical disparities in sampling schemes is that we are still lacking a common and unifying framework to compare large scale species diversity patterns between groups of the marine plankton. Such a framework would enable us to test hypotheses about the relative importance of the underlying drivers of species diversity and to investigate its link with ecosystem functions such as productivity, resource use and the efficiency of export production (Biodiversity-Ecosystem Functioning relationships; Cardinale et al., 2012). On land, the current paradigm is that ecosystem functions and their temporal stability show a positive and saturating relationship with increasing species diversity as a result of the selection and complementarity processes (Cardinale et al., 2012; Gonzalez et al., 2020). Through the asynchronous selection of those species that are the fittest, more diverse communities tend to display a range of species capable of exploiting the various resources available for converting them into biomass. This paradigm mainly emerged from local experiments on land where relatively few species were artificially manipulated to examine the relationship between species richness and functions such as productivity or resource use (Cardinale et al., 2012; Tilman et al., 2014). It is still unclear how this paradigm holds against field observations of marine plankton (Gamfeldt et al., 2015). Very few studies documented Biodiversity-Ecosystem Functioning relationships for phytoplankton communities and they usually reported conflicting results (Ptacnik et al., 2008; Cermeño et al., 2013; Vallina et al., 2014; Lehtinen et al., 2017; Acevedo-Trejos et al., 2018; Chen et al., 2019). Even fewer studies investigated Biodiversity-Ecosystem Functioning relationships for marine zooplankton (Beaugrand et al., 2010; Guidi et al., 2016; Hébert et al., 2016). Consequently, on top of lacking fundamental knowledge on the contemporary global diversity gradients of various plankton groups, we remain even further away from understanding how such diversity gradients relate to the performance of ecosystem functions like nutrient use efficiency, productivity and carbon export.

Here, we use the recent compilations of plankton species occurrences and ensemble species distribution models (SDMs) framework of Benedetti et al. (2021) to: (i) hierarchize the environmental predictors of species distributions that are then used to (ii) estimate the global contemporary richness patterns for 14 different functional group (instead of broad trophic levels) and evaluate how they covary with key ecosystem properties (i.e., net primary production, carbon export efficiency or higher trophic level diversity). By doing so, we aim to provide spatially explicit information about plankton biodiversity to the scientific community so it can explore emerging patterns in Biodiversity-Ecosystem Functioning relationships and put forward hypotheses that can be tested by future studies. We also hope to shed more light on the diversity patterns of plankton groups that have been historically understudied.

## 2. Methods

### 2.1. Plankton occurrence data

Species-level plankton occurrences (presence-only) were retrieved from two recent efforts aiming to describe and model phyto- and zooplankton communities in the global open ocean (Righetti et al., 2019; Benedetti et al., 2021). Plankton are commonly divided between two trophic levels: the photoautotrophic phytoplankton and the heterotrophic zooplankton. Both categories comprise a tremendous variety of species belonging to various clades and are characterized by different biological requirements and thus niche dimensions. The steps described below, from data acquisition to SDMs parameterization, reflect this major initial dichotomy.

The phytoplankton species occurrence data used here were compiled by Righetti et al. (2020) and stem from various sources: Global Biodiversity Information Facility (GBIF; https://www.gbif.org), Ocean Biogeographic Information System (OBIS; https://www.obis.org), Villar et al. (2015), and MAREDAT (Buitenhuis et al. 2013). The dataset gathers >10^6^ occurrences for nearly 1700 species sampled through diverse techniques within the monthly climatological mixed layer depth, at an average depth of 5.4 ± 6.9 m (mean ± sd) for the 1800-2015 period. Phytoplankton species names were corrected and harmonized following the reference list of Algaebase (http://www.algaebase.org/). The resulting species list spans most of the extant phytoplankton taxa contributing to the diversity of the phytoplankton communities prevailing in the euphotic zone of the oceans (Righetti et al. 2020). This dataset has been used to derive a surface open ocean estimate of the global phytoplankton diversity that properly accounts for sampling spatial-temporal biases and that was validated against independent data (Righetti et al., 2019; Benedetti et al. 2021).

The zooplankton species occurrences used here were compiled by Benedetti et al. (2021). In short, occurrences were primarily retrieved from OBIS and GBIF and complemented by the observations from Cornils et al. (2018) and Bednaršek et al. (2012). The data cover all main extant marine zooplankton clades. Occurrences that are not representative of contemporary open ocean plankton communities were discarded based on the following criteria: (i) a missing spatial coordinate, (ii) incomplete sampling date (d/m/y), (iii) year of collection older than 1800, (iv) no sampling depth provided or sampling depth > 500 m (i.e. excluding taxa mainly inhabiting the deep ocean while accounting for those performing vertical migrations), (v) not identified to the taxonomic species level, (vi) issued from drilling holes or sediment samples, (vii) associated with monthly surface salinity values < 20, and (viii) collected in a 1°x1° grid cell displaying a seafloor shallower than 200m (i.e. excluding coastal areas). Zooplankton species name were carefully harmonized following the reference list of the World Register of Marine Species (WoRMS; http://www.marinespecies.org). The final zooplankton dataset compiled occurrences for > 3000 different taxa spanning the main zooplankton clades and functional groups.

### 2.2. Plankton functional groups

The definition of plankton functional groups (PFGs) depends on the scientific question addressed (Le Quéré et al., 2005; Hood et al., 2006). Most of the time, taxa are grouped into PFGs based on the performance of particular biogeochemical functions (e.g. calcification vs. silicification; Iglesias-Rodriguez et al., 2002) and/or size classes that describe their trophic level in food-webs and how they contribute to pathways of carbon fluxes (e.g. nano-, micro-, or mesoplankton). Here, we focus on 14 PFGs (three for phytoplankton and eleven for zooplankton) displaying at least 5 different species whose distributions could be modelled from the available occurrences data (see section 2.3 below). The PFGs investigated and their main functions are described in Table 1. These PFGs were made to include taxa from either the same size class and/or that are known to fill the same trophic niche as a result of similar feeding traits (Kiørboe, 2011). Because the observational datasets do not cover the smallest size classes of the plankton (Righetti et al., 2020; Benedetti et al., 2021), some key functional groups (e.g., N2-fixing prokaryotes, picophytoplankton, microzooplankton) could not be integrated in this study.

**Table 1:**
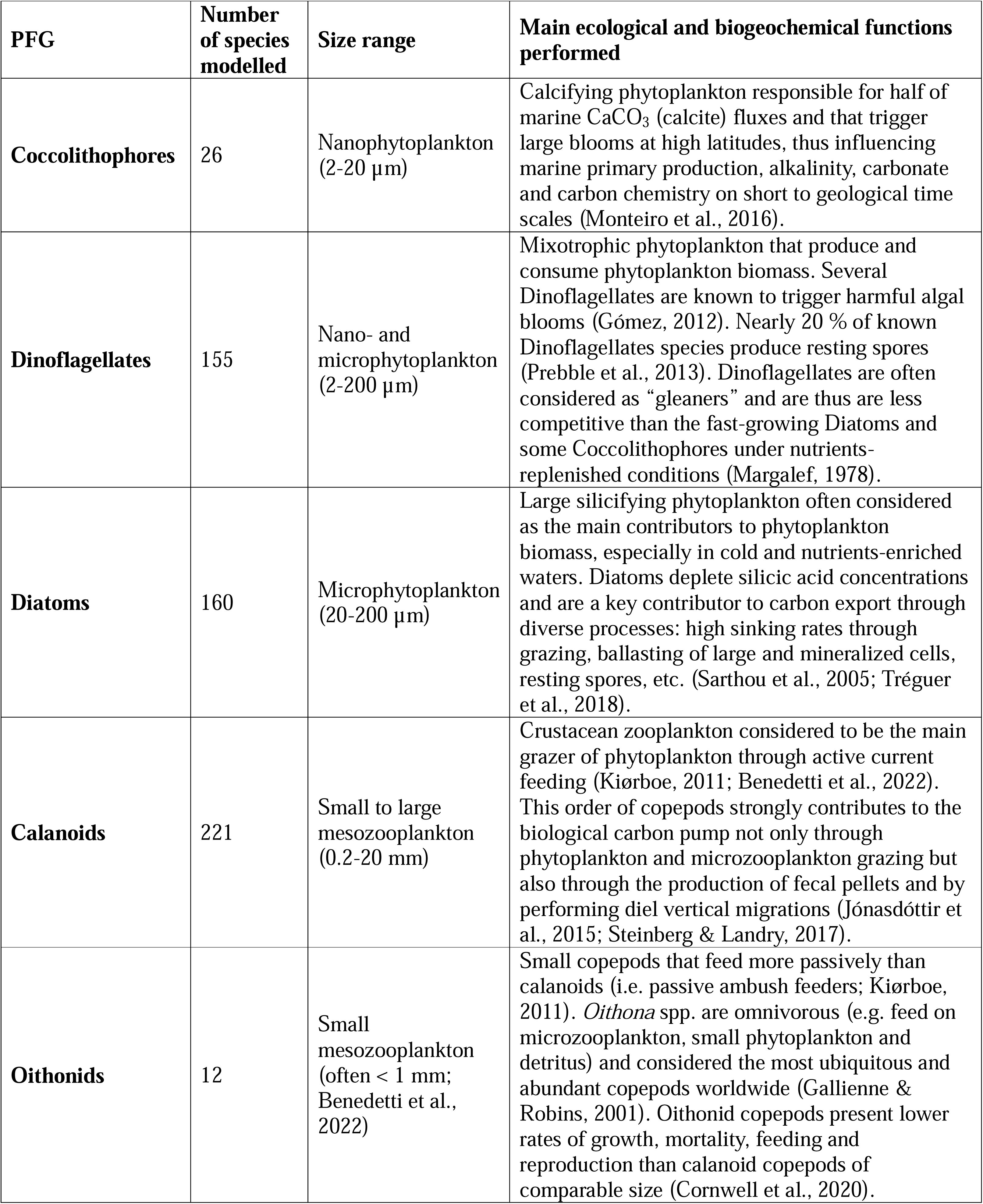
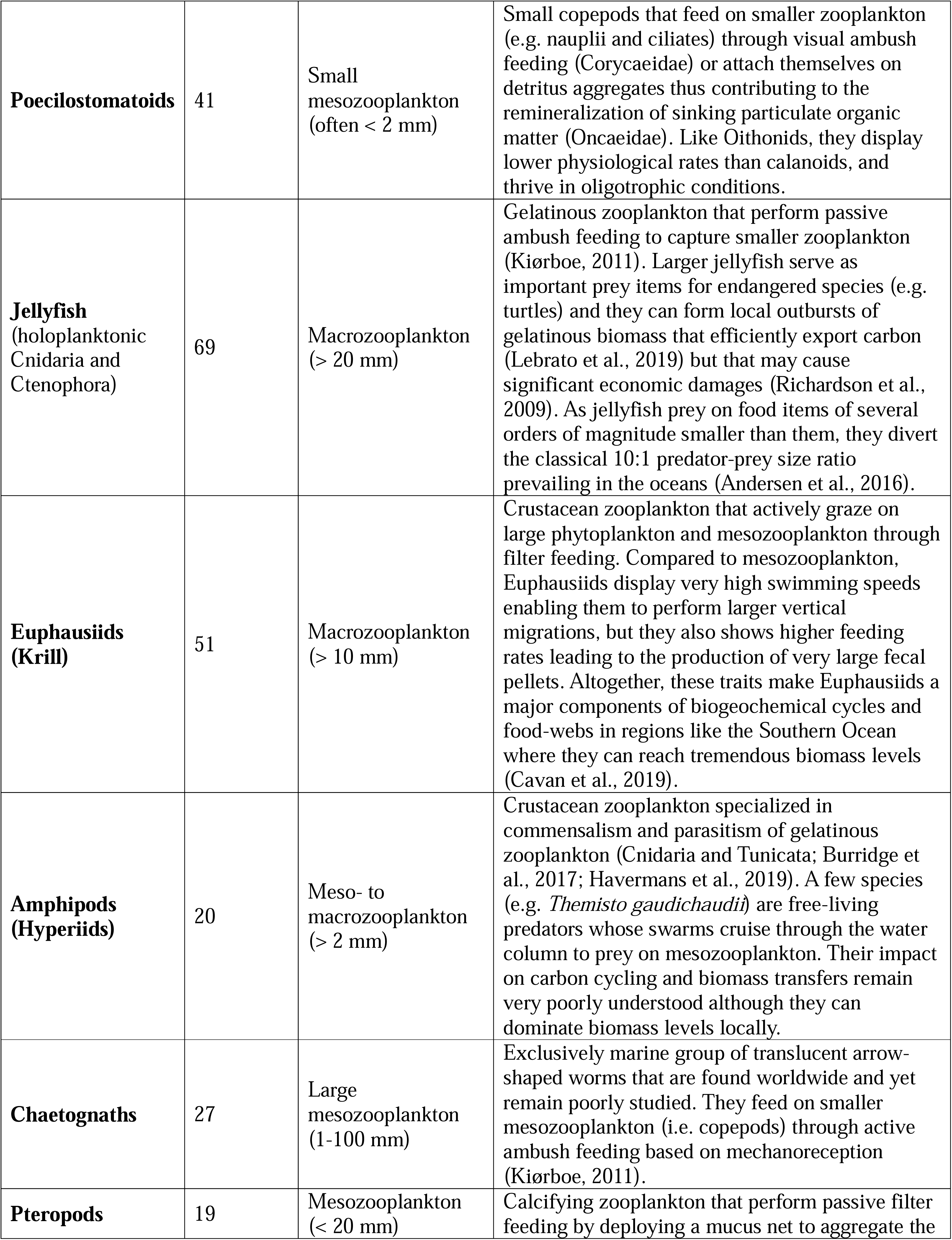
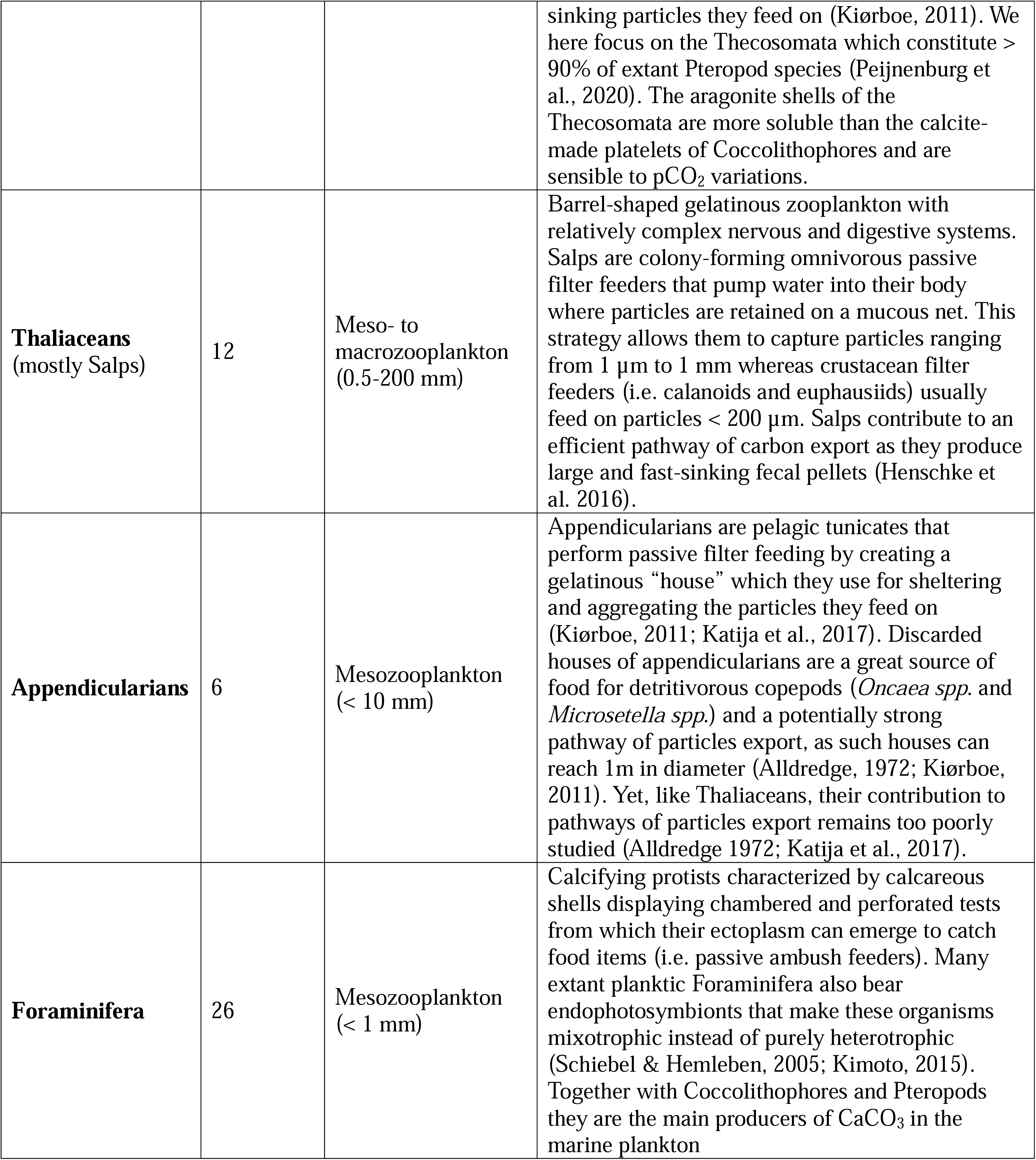
Overview and description of the 14 plankton functional groups (PFGs) whose species diversity is modelled in the present study.

In total, 845 different species could be classified into one of the above PFGs and were considered for modelling. Diatoms, Dinoflagellates and Coccolithophores constitute 47%, 45% and 8% of the phytoplankton species, respectively. Calanoids represent 44% of all zooplankton species and all other zooplankton groups show relative contributions to total species richness of < 15% (Figure S1; Table S1).

### 2.3. Species distribution modelling

#### 2.3.1. Model types

SDMs were developed following the standard ensemble modelling approach of Benedetti et al. (2021) which covers a range of model types and complexity: Generalized Linear Models (GLM), Generalized Additive Models (GAM) and Artificial Neural Networks (ANN). To prevent the pitfalls associated with model over-fitting (Merow et al., 2014), we restricted the number of environmental predictors relative to the number of presences (see below) and SDMs were tuned to fit relatively simple response curves. The full details of the model parametrization are given in Benedetti et al. (2021). All SDMs were developed using the ‘*biomod2’* R package (Thuiller et al., 2020).

#### 2.3.2. Background data

Background data were drawn to inform the SDMs as to which environmental conditions were sampled by the observations but where a species is not likely to occur (i.e., pseudo-absences; Barbet-Massin et al., 2012). We used of the target-group approach of Philipps et al. (2009) which draws background data as a function of the presence data distribution and thus does not induce additional biases and avoids misclassifying unsampled regions as unsuitable habitats. The PFGs were used as target-groups, meaning that the background data of each species were randomly drawn from the sites (i.e. monthly 1°x1° grid cells) displaying at least one occurrence of their corresponding PFG, while omitting those sites where the species was found as present. Four zooplankton PFGs displayed too sparse occurrences to be considered as target-groups (Amphipods, Thaliaceans, Appendicularians and Foraminifera; Figure S2). In this case, the background data of the corresponding species were randomly drawn from the total pool of sites with zooplankton occurrences (i.e. total background approach; Righetti et al., 2023). Here, only those sites presenting at least one occurrence from five different PFGs were considered to avoid including sites where oceanographic cruises focused on a small subset of the plankton community (e.g., only diatoms or only pteropods). For each species, 10 times more background data than presences and background data were weighted inversely proportional to the presences (Barbet-Massin et al., 2012). Ultimately, all background data were matched with monthly values for the predictors considered environmental predictors for the SDMs.

#### 2.3.3. Selection of environmental predictors

A comprehensive set of monthly climatologies for 20 environmental predictors that affect the physiology and the distribution of plankton species was implemented onto the standard 1°x1° global cell grid of the World Ocean Atlas (WOA; see Table S2 for exhaustive references). First, eleven primary predictors were retrieved: sea surface temperature (SST, °C); surface concentration of nitrate (NO_3_^-^), phosphate (PO_4_^3-^) and silicic acid (Si(OH)_4_) (µmol L^-1^); dissolved oxygen concentration at 175m depth (O_2_, mL L^-1^); photosynthetically active radiation (PAR; µmol m^−2^ s^−1^); surface phytoplankton chlorophyll-a (Chl-a, mg m^-3^); mixed-layer depth (MLD, m); surface wind stress (Wind, m s^−1^); surface partial pressure of carbon dioxide (pCO_2_, µatm); and eddy kinetic energy (EKE, m^2^ s^-2^) as a proxy of the strength of mesoscale activity. Then, nine secondary predictors were derived from the eleven primary ones: PAR over the MLD (MLPAR, µmol m^−2^ s^−1^); excess of NO_3_^-^ to PO_4_^3-^ relative to the Redfield ratio (N* = NO_3_^-^ - 16*PO_4_^3-^; Gruber & Sarmiento, 1997); and excess of Si(OH)_4_ relative to NO_3_^-^ (Si* = Si(OH)_4_ – NO_3_^-^; Sarmiento et al., 2004). Since the distribution of macronutrients concentrations, Chl-a and EKE were highly skewed towards low values, we examined their logarithmic transformations (log(NO_3_), log(PO_4_), log(Si(OH)_4_), log(EKE), and log(Chl-a) (based on the natural log) as additional secondary predictors as these were closer to a normal distribution.

Collinearity between predictors can inflate model parameters, mislead importance rankings in regression models and thus bias SDMs projections (Dormann et al., 2013). Therefore, we investigated predictor collinearity by computing pairwise Spearman’s rank correlation coefficients (?) from each species-specific dataset. Median pairwise correlation coefficients were derived from the distribution of species-levelvalues and one of two predictors from a pair displaying a median |?| > 0.75 was discarded (Dormann et al., 2013). Preference was given to those predictors closest to a normal distribution and least correlated to SST.

Several criteria were used to select the predictors for the three phytoplankton groups from the initial set. O_2_ was excluded as a potential predictor for phytoplankton PFGs owing to their photoautotrophic nature. Nutrient predictors (except N* and Si*) tend to be strongly correlated with several pairs exhibiting median |?| > 0.75. Consequently, NO_3_^-^ and PO_4_^3-^ concentrations were discarded and only log(Si(OH)_4_) was retained as a predictor representing the global gradient in nutrient availability as its medianwith SST is lower (median= -0.15) than that for NO_3_^-^ and PO_4_^3-^ (median= -0.51 and -0.17, respectively). Plus, Si(OH)_4_ is a limiting nutrient for shaping diatom growth and distributions. We used its log-transform since it showed a more normal distribution than Si(OH)_4_. Log(PO_4_) has a medianwith N* of -0.82 so N* was kept. The other cluster of highly correlated predictors comprised MLD, PAR and MLPAR (median= -0.78 and 0.84 between MLPAR and MLD and then PAR, respectively). MLPAR was discarded as the medianbetween MLD and PAR was -0.69, and MLD was very weakly correlated with SST (median= -0.07). The final ten predictors retained for phytoplankton PFGs were: SST, PAR, log(Chl-a), Wind, MLD, N*, Si*, log(Si(OH)_4_), log(EKE) and pCO_2_.

For zooplankton groups, the same criteria were used to select their subset of predictors. The predictors at the location of the observations display similar patterns in pairwise median correlationsas seen for phytoplankton, except for the macronutrients that have even stronger correlations (all pairwise median> 0.80) and SST (all pairwise median< -0.75 except for log(Si(OH)_4_)). Therefore, the same predictors as for phytoplankton were retained for the zooplankton PFGs, with the addition of O_2_.

#### 2.3.4. Ranking of predictors importance across PFGs

To further narrow down the number of PFG-specific predictors to be included in the SDMs, univariate permutation tests were performed to rank the retained predictors. For every species-specific dataset, one of the predictors was randomly reshuffled and a multivariate SDM was trained and its projections was compared to those from the non-reshuffled SDM through a Pearson’s correlation coefficient. The latter was then subtracted from 1, thus returning a quantitative score varying between 0 and 1 which indicates the importance of the reshuffled covariate for the performance of the multivariate SDM and thus for constraining the modelled species’ distribution. For each species and SDM type, 30 univariate random permutation experiments were carried out, which lead to 90 scores of relative importance per predictor.

Non parametric variance analysis (Kruskal-Wallis rank sum tests) was performed to assess the variations of each predictor’s ranking across all species modelled, phyto- and zooplankton species separately, and across PFGs. Dunn’s pairwise multiple comparison rank sum tests were then applied with a Bonferroni correction to identify pairs of predictors displaying significant variations in rankings.

For each PFG, the six predictors displaying the highest scores were retained to model the distribution of the species belonging to the corresponding PFG. We retained six predictors as all the species modelled present > 75 occurrences and because we aimed to achieve a presence-to-predictors ratio > 12:1, which is higher than the 10:1 ratio recommended by Guisan et al. (2017). Concomitantly, each SDM was run 10 times with a five-fold cross validation and the resulting True Skill Statistics (TSS; Allouche et al., 2006) was calculated to evaluate SDMs skills. Once the six PFG-specific predictors were selected, the species presences-background data were randomly split into ten different training and testing sets (80%-20% respectively). Therefore, 30 models (3 SDMs types x 10-fold cross validation) were trained per species. Model skill was evaluated through the TSS. All 845 species displayed an average TSS > 0.30 so they were all retained.

#### 2.3.5. Projections of mean annual species richness

For each species, the 30 models were projected onto the twelve monthly climatologies of the predictors included in the SDMs, thus generating global projections of monthly habitat suitability indices (HSI). The latter highlight the regions where environmental conditions are most favorable for a species to be present. Since the main goal of our study is to study species richness patterns, each model’s HSI projection was converted to a presence-absence distribution map based on the HSI threshold that maximized the TSS score of the model. For each PFG, the model-specific monthly maps were stacked to estimate their monthly richness and species composition. Mean annual richness was then derived by averaging monthly estimates for each model. Following an ensemble projection approach, the final ensemble projections of mean annual richness were obtained by averaging the model-specific annual richness estimates. True richness levels cannot be estimated here since many taxa that contribute to the richness of the present PFGs in nature cannot be modelled due to the limited occurrence density. Therefore, the final species richness estimates were divided by the number of species modelled in each PFG to present species diversity estimates that represent the emergent global gradients in PFGs diversity.

Because the species richness estimates were normalized to the number of species modelled, they range between 0 and 1 with values equal to 1 indicating the grid cells where environmental conditions allow all species to be modelled as present (i.e., species accumulation). Based on these estimates, we explored the inter-PFGs differences in latitudinal diversity gradients in two ways. First, we quantified the strength of the covariance between mean latitudinal richness and absolute latitude through rank correlation coefficients (?). This way, we identified which PFGs show a diversity gradient that can be predicted by latitude, whatever the strength of this gradient. Second, we fitted linear regressions between mean latitudinal species richness and absolute latitude. The linear regressions were not computed across the full latitudinal range as most PFGs displayed non monotonic and nonlinear latitudinal gradients of richness (Document S1). Here, we aimed to compare PFGs on the basis of their relative rate of species accumulation, rather than trying to prove the existence of linearity among the latitudinal gradients. To do so, we computed the first derivatives of the mean latitudinal annual richness over absolute latitude to detect the latitudinal ranges over which the groups’ richness shows a monotonic and nearly linear decrease in species richness (Document S1). Absolute latitudes > 65° were excluded since changes in diversity beyond this threshold only occur within a very small (often < 0.1) ranges of species richness. The linear regressions were fitted from the first absolute latitude that is followed by ten consecutive negative derivative values (i.e., a monotonic decrease in richness) to the first latitude showing a positive derivative (i.e., richness increase). Two PFGs (Chaetognaths and Foraminifera; Document S1) showed a local maximum (i.e., ten consecutive positive derivative values) between two phases of richness decrease. We chose to integrate these local maxima into the linear regressions instead of fitting two smaller and separate linear models for the same PFG. The slopes of the linear regressions were then used to compare the steepness of the latitudinal diversity gradient (i.e., rate of species accumulation over latitude) between PFGs through covariance analysis (ANCOVA).

### 2.4. Analyses

#### 2.4.1. Similarity between richness patterns and their covariance with ecosystem properties

To evaluate the similarity between the patterns of annual richness across PFGs and help delineate their main modes of spatial variability, a principal component analysis (PCA; Legendre & Legendre, 2012) was performed on the ensemble estimates of mean annual richness. PCA is a dimensionality reduction analysis that summarizes the correlation structure between input variables through a symmetric covariance matrix whose ensuing eigenvectors are used to construct principal components (PCs). PCs are linear combinations of the input variables that are orthogonal to each other so they summarize uncorrelated modes of spatial variability in richness, and are ranked according to their relative contribution to explained variance.

We aimed to determine the main environmental covariates of species richness, so the annual climatologies of the following environmental predictors were added as supplementary variables in the PCA: SST, O_2_ (at 175m depth), PAR, log(Chl-a), Wind, MLD, log(NO3), log(Si(OH)_4_), N*, Si*, log(EKE) and pCO_2_. Although they were not considered as candidate predictors for the SDMs, mean annual sea surface salinity (SSS) and the annual range of SST (dSST; maximum monthly value - minimum monthly value) were added in the PCA as supplementary variables to better represent the range of environmental conditions experienced by PFGs in the ocean. This way we can examine the covariance between the PFGs-level richness and environmental predictors. We highlight that this is not a truly independent test as the monthly climatologies of some of these variables were used as predictors in the SDMs (see section 2.3.3).

Furthermore, we aimed to examine the covariance between PFG species richness and five proxy variables of ecosystem functioning to investigate emergent Biodiversity-Ecosystem Functioning relationships. The following five variables represent plankton-related processes or factors that provide key socio-economical services: (i) normalized global species richness of oceanic taxa (i.e., bony fishes, sharks, cetaceans and squids) which is indicative of overall marine megafauna biodiversity (Tittensor et al., 2010); (ii) mean annual surface net primary production (NPP, mg Carbon m^-2^ day^-1^) (DeVries & Weber, 2017); (iii) the corresponding flux of particulate organic carbon (FPOC, mg Carbon m^-2^ day^-1^) that is exported below the euphotic zone which indicates the strength of the biological carbon pump (DeVries & Weber, 2017); (iv) the FPOC/NPP ratio which indicates the efficiency of the biological carbon pump; and (v) the inverse of the mean annual slope of the power-law particles size distribution measured from satellites (Kostadinov et al., 2009), which serves as a quantitative particle size index (PSI; the higher the value, the higher the contribution of larger particles and phytoplankton cells to the total size spectrum). These five proxies of marine ecosystem services were also included in the same PCA as supplementary variables. This way, this single multivariate analysis enabled us to: (i) identify the PFGs that display similar mean annual richness patterns; (ii) identify the main environmental gradients covarying with these species richness patterns; and (iii) explore the emerging patterns between the richness of various PFGs and proxy variables of ecosystem functioning.

#### 2.4.2. Uncertainties in projections of mean annual PFGs species richness

The present global richness estimates are sensitive to sources of uncertainty, i.e. factors of the modelling framework, such as SDM type choice or species prevalence, that lead to inter-model variability in species richness projections (Thuiller et al., 2019). Evaluating the uncertainty levels can help identify the PFGs for which current SDMs approaches provide less robust richness estimates. The standard deviation of the 30 model-specific annual richness projections was computed for each PFG to quantify their relative level of uncertainty in annual richness estimates. The linear relationship between this estimate of relative uncertainty and the number of species modelled and/or sampling effort per PFG (i.e., the number of occurrences available for training the SDMs) was examined to test if uncertainties in species richness estimates decrease with higher sampling effort.

## 3. Results

### 3.1 Relative importance of environmental predictors for constraining species distributions

Overall, SST emerged as the top-ranking predictor constraining the species distributions (Figure 1; median rank ± IQR = 0.42 ± 0.51), phytoplankton groups (0.35 ± 0.45) and zooplankton groups (0.47 ± 0.53). The rankings of the other predictors depended on the aggregation level. When taking all 851 plankton species together, SST was followed by: O_2_ (zooplankton only), N*, Si*/log(Si(OH)_4_) (non-significant variations based on Dunn’s pairwise multiple comparison tests), Wind/PAR, MLD, log(EKE)/log(Chl-a), and pCO_2_ was found the least important variable. For phytoplankton species, SST was followed by: N*, Wind, PAR/Si*/log(Si(OH)_4_), and the pCO_2_/MLD,/log(Chl-a)/log(EKE) group. For zooplankton species, SST was followed by: O_2_/Si*/log(Si(OH)_4_), PAR/Wind/MLD, log(EKE)/N*, log(Chl-a) and then pCO_2_ again as the least performing predictor. SST remained the top predictor at the PFG-level, except for coccolithophores and pteropods whose top ranking predictors were N* and O_2_, respectively (Figure 1c,j). The top six ranking variables of each PFG was retained to train the SDMs are given in Table S3.

**Figure 1:**
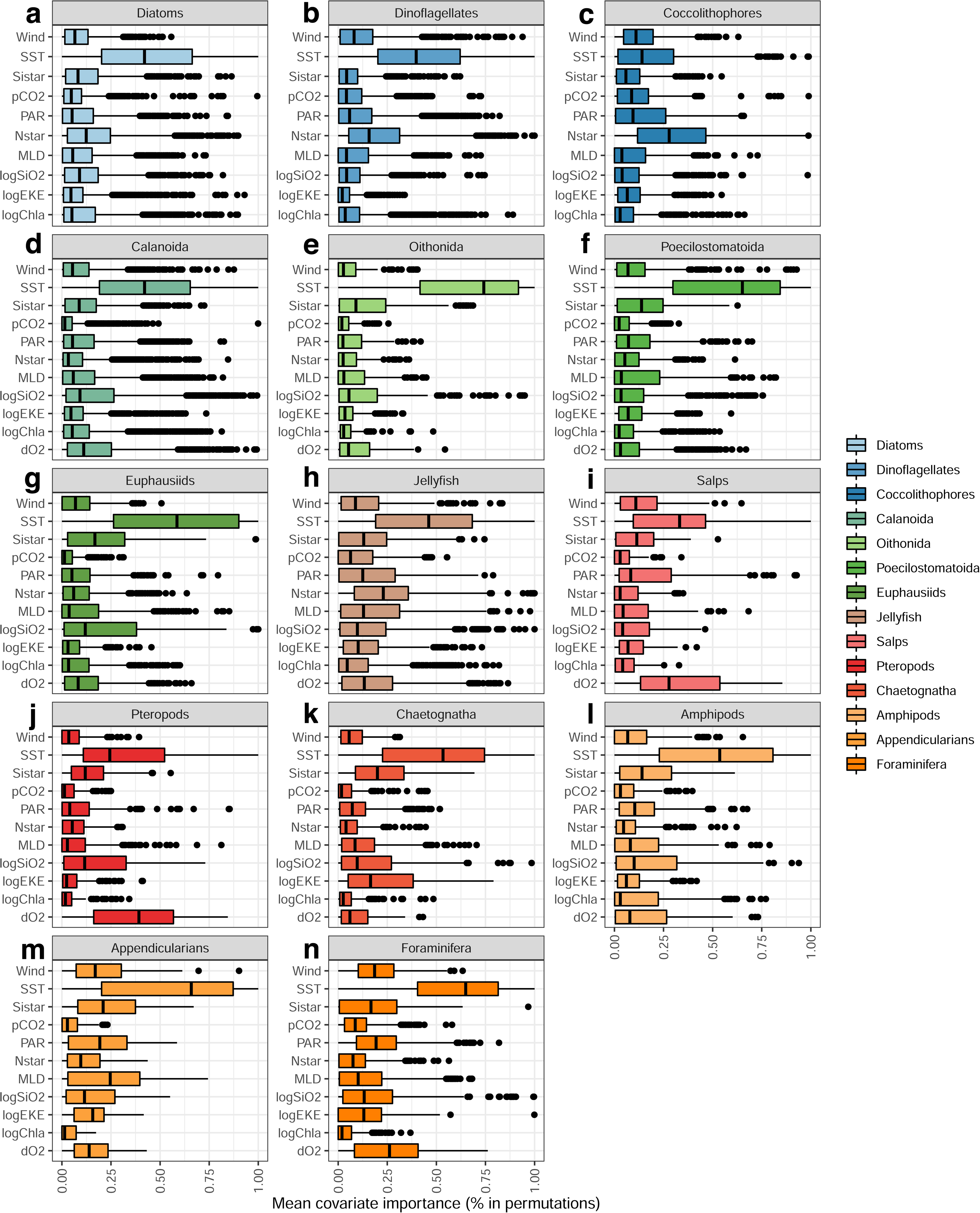
Distribution of the ranks of relative importance of each environmental predictor investigated for modelling the species distributions of a) Diatoms (n = 163), b) Dinoflagellates (n = 157), c) Coccolithophores (n = 27), d) Calanoida (n = 221), e) Oithonida (n = 12), f) Poecilostomatoida (n = 41), g) Euphausiids (n = 51), h) Jellyfish (n = 69), i) Salps (n = 12), j) Pteropods (n = 19), k) Chaetognatha (n = 27), l) Amphipods (n = 20), m) Appendicularians (n = 6), and n) Foraminifera (n = 26). For each predictor and species, ranks were determined through 30 random permutation tests for each species distribution model (30 models, so n = 90 scores in total). Ranks were then aggregated at the plankton functional group level. The central vertical lines indicate the median values, the boxes illustrate the interquartile ranges and the error bars indicate 5^th^ and 95^th^ percentiles.

### 3.2 Global latitudinal gradients in mean annual species richness

All PFGs display decreasing species richness from the low to the high latitudes and all showed peaks in mean annual richness within the tropical band (0-30°) or near it (Figures 2 & 3). Yet, PFGs show marked contrasts in their latitudinal diversity gradients as their richness showed varying strength in covariance with absolute latitude and varying rates of species accumulation (Figure 2). Most groups showed strong negative correlations with absolute latitude, withvalues ranging between -0.70 and -0.80 (all p < 0.001). Coccolithophores richness shows the weakest covariance with latitude (? = -0.52) while Dinoflagellates richness shows the strongest (? = -0.93). ANCOVA reveals that the coefficients of the linear regressions fitted vary significantly between PFGs (F = 1256.11; p < 0.001), indicating that most PFGs show different rates of decrease in species accumulation along the monotonic parts of the richness gradients. Posthoc Tukey’s multi-comparison tests confirm that most pairs of PFGs show significant differences in the slope of the linear models (adjusted p < 0.001; Document S1). Overall, Oithonids, Pteropods, Poecilostomatoids, and Salps show the steepest latitudinal diversity gradients (slopes < -0.02). Meanwhile, Amphipods, Appendicularians, Chaetognaths, Diatoms, Dinoflagellates and Coccolithophores show the weakest latitudinal diversity gradients (slopes > -0.004). Phytoplankton groups tend to show weaker slopes and therefore weaker rates of decrease in richness than most of zooplankton groups. Among the zooplankton, Pteropods, Salps, Jellyfish and the three copepod groups show the strongest weaker rates of decrease in richness from the tropics to the high latitudes. In the tropical band, all PFGs show mean annual richness values above or close to 0.40 except Amphipods (mean ± sd = 0.28 ± 0.02). In the polar areas (> 60°), all PFGs show mean diversity values > 0.10 except the Calanoids, Poecilostomatoids, Euphausiids and Salps. The spatially explicit patterns of global PFGs richness are given in Figure 3 to illustrate the longitudinal variability (i.e., light blue ribbons on Figure 2) on top of the main latitudinal patterns. Correlation coefficients were computed between species richness and longitude (spanning 0-360°) to test for potential longitudinal patterns as well. All PFGs display significant (all p < 0.01) but quite weak (all |?| < 0.2) longitudinal patterns of richness. Since these correlation coefficients are much weaker than those based on latitude (Figure 2) we choose not to comment them further. The maps also highlight the relative position of the peaks and dips in mean annual richness of each functional group (but see section 3.3 right below for further description).

**Figure 2:**
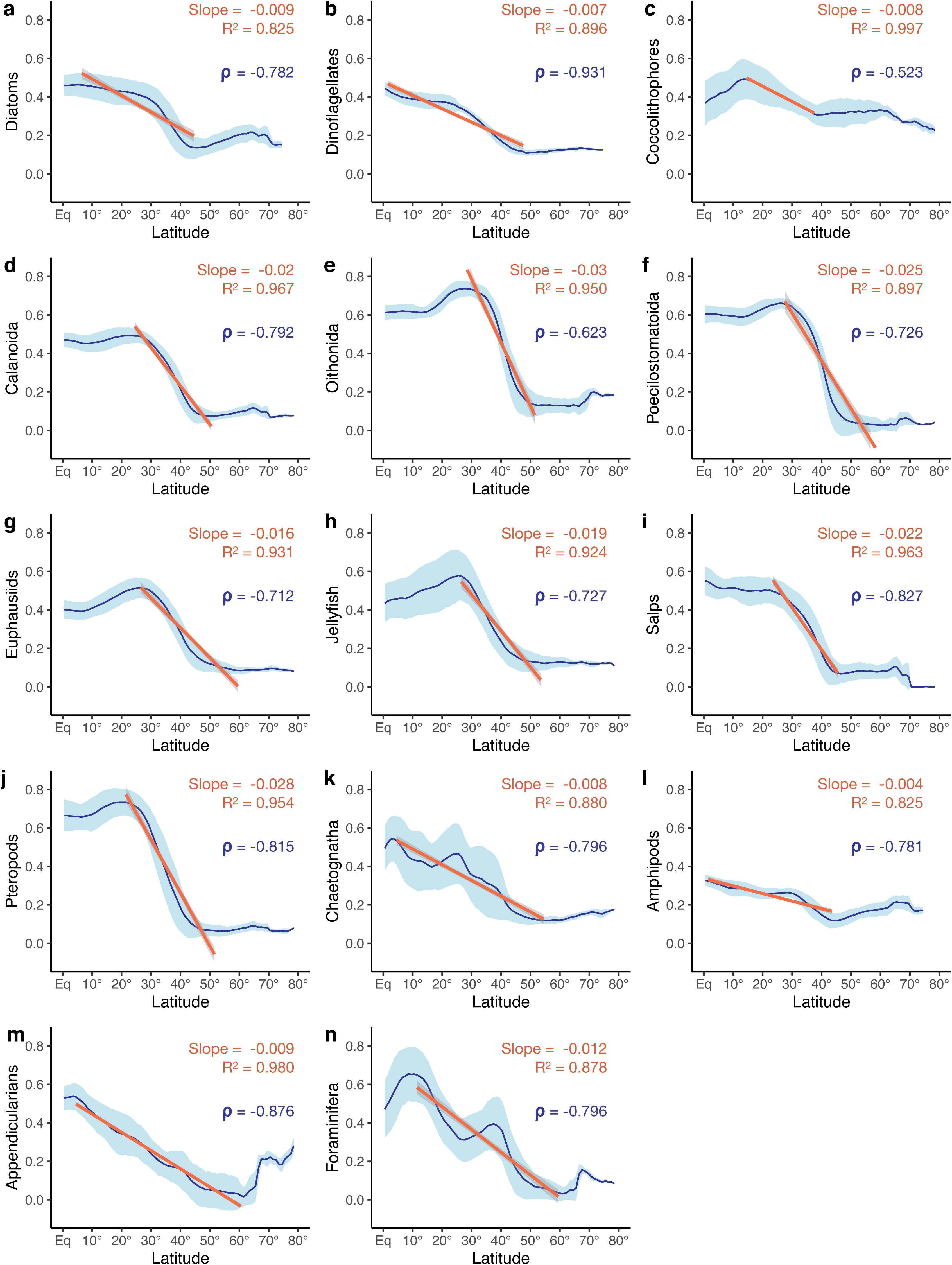
Zonal distribution of mean annual richness (expressed in % of species modelled) as a function of absolute latitude for a) Diatoms (n = 163), b) Dinoflagellates (n = 157), c) Coccolithophores (n = 27), d) Calanoida (n = 221), e) Oithonida (n = 12), f) Poecilostomatoida (n = 41), g) Euphausiids (n = 51), h) Jellyfish (n = 69), i) Salps (n = 12), j) Pteropods (n = 19), k) Chaetognatha (n = 27), l) Amphipods (n = 20), m) Appendicularians (n = 6), and n) Foraminifera (n = 26). The blue curves illustrate the latitudinal averages of species richness per absolute degrees of latitude. The light blue ribbons illustrate the standard deviation associated with the latitudinal averages (i.e., longitudinal variability). The solid orange lines illustrate the linear regressions computed over the latitudinal range of interest. The slope and the adjusted R^2^ of the linear regressions indicate the strength of the decrease in species accumulation. The Spearman’s correlation coefficients (?) indicate the strength of the covariance between absolute latitude and average species richness. All model coefficients and correlation coefficients show p-values < 0.001.

**Figure 3:**
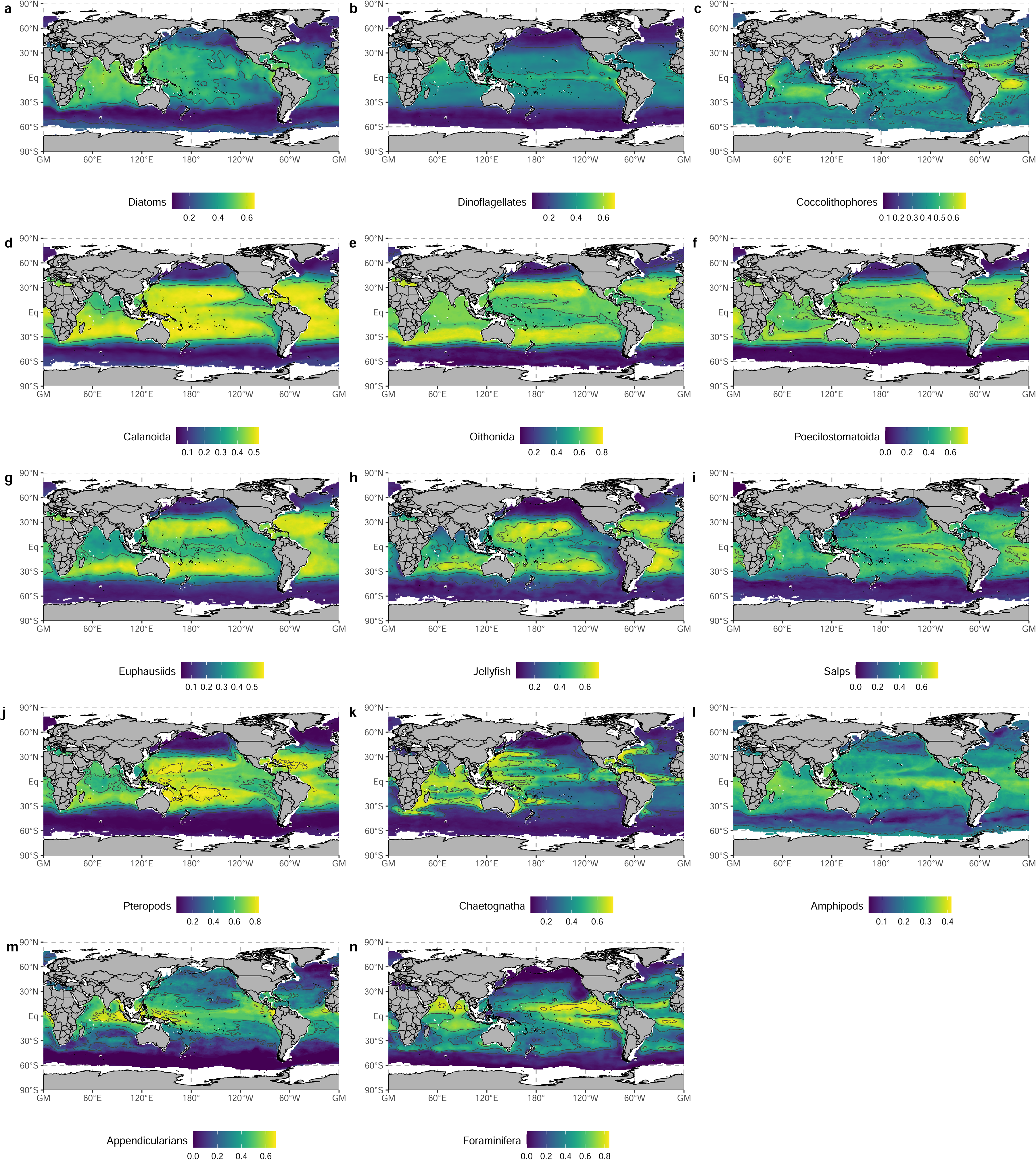
Spatial distribution of the mean annual species richness (expressed in % of species modelled) for a) Diatoms (n = 163), b) Dinoflagellates (n = 157), c) Coccolithophores (n = 27), d) Calanoida (n = 221), e) Oithonida (n = 12), f) Poecilostomatoida (n = 41), g) Euphausiids (n = 51), h) Jellyfish (n = 69), i) Salps (n = 12), j) Pteropods (n = 19), k) Chaetognatha (n = 27), l) Amphipods (n = 20), m) Appendicularians (n = 6), and n) Foraminifera (n = 26). The data used here are the same as in Figure 2. Blank areas correspond to those grid cells where richness projections were not possible for more than six months due to missing values in the monthly environmental climatologies the models were projected on.

### 3.3 Emergent relationships between species richness and environmental covariates

A PCA identified the equatorward increase in richness to be the first-order pattern as PC1 summarizes nearly 77% of the spatial variance and all PFGs scored positively along PC1 (Figure 4a,c). While PC2 explains a lower amount of variance (7.5%; Figure 4b) it allows us to distinguish PFGs based on the location of their relative peaks and troughs in mean annual richness, permitting us to investigate productivity-related patterns that are decoupled from the global temperature gradient. PC2 is mainly scored by the species richness of Coccolithophores, Jellyfish and Euphausiids on the positive side and Appendicularians, Amphipods, Diatoms and Dinoflagellates on the negative side (Figure 4d). Consequently, the regions in blue on Figure 4b correspond to those regions where the latter four PFGs have peaks in species richness, whereas the regions in red indicate those where the richness of Coccolithophores, Jellyfish, Euphausiids peak.

**Figure 4:**
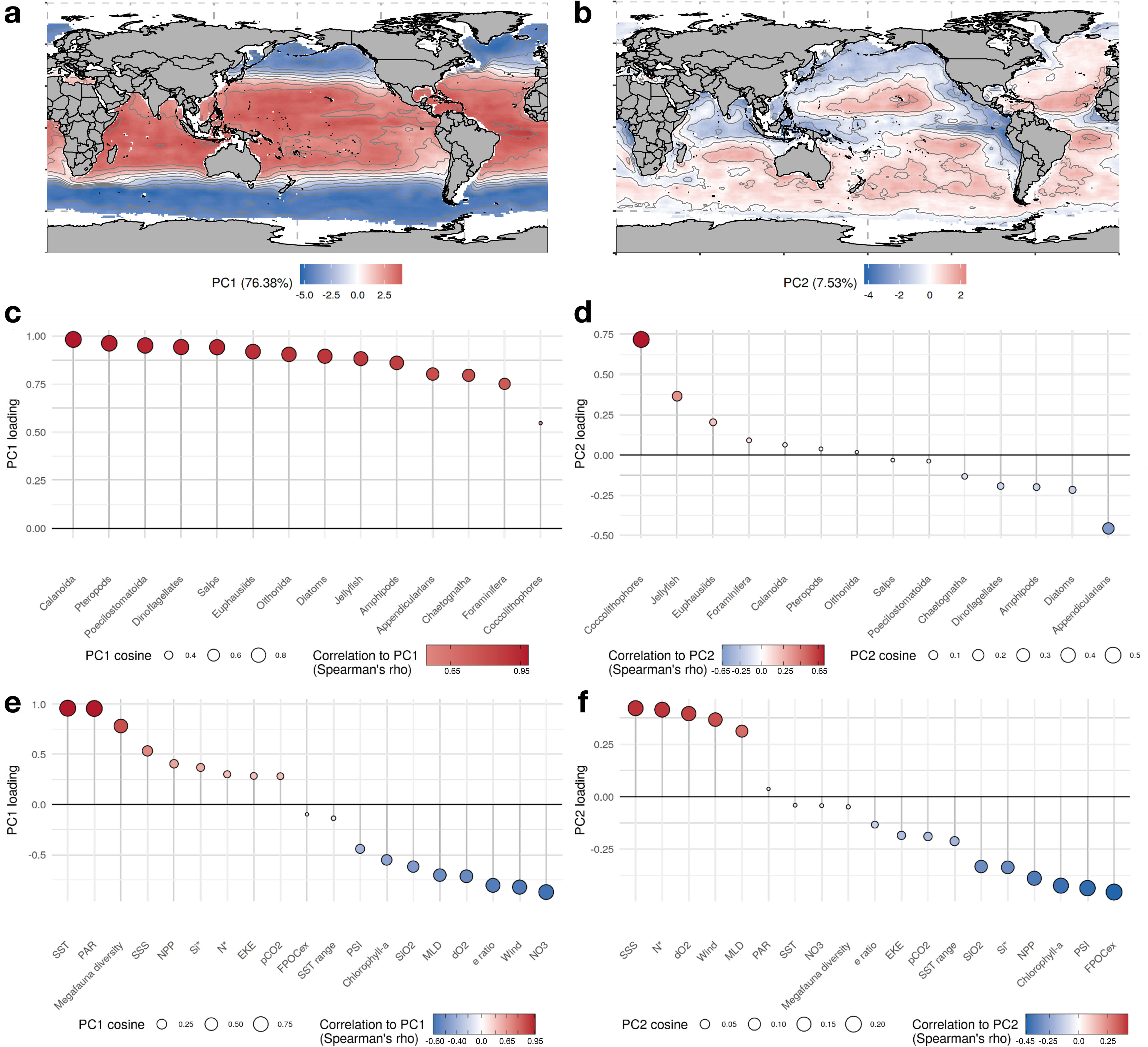
Projections of the first two components of a principal component analysis (PCA) based on the ensemble projections of mean annual species richness of the plankton functional groups for the global surface open ocean (n = 29680 ocean grid cells). Maps a) and b) show the spatial variability of PC1, PC2 respectively. The lollipop plots in c) and d) indicate the loadings of the groups’ annual species richness that were used to construct PC1 and PC2. The lollipop plots in e) and f) indicate the loadings of the mean annual climatologies of environmental predictors that were added as supplementary variables in the PCA to analyze their covariance with PC1 and PC2. In the lollipop plots, the circles are colored as a function of the variables’ correlation to the PCs. Circles in red correspond to those variables that covary positively with the regions in red on the maps (and vice versa for the variables in blue). The size of the circles indicates the quality of the variables’ projection in the PCA space based on their squared cosine.

The PCA also allows us to uncover the combination of environmental variables that covary most strongly with the patterns of species richness, including those predictors that were not used in the SDMs (Figure S4). The environmental covariates that display a strong latitudinal gradient on a mean annual scale display the strongest covariation with PC1: SST and PAR with positive correlations, and log(NO3), log(Si(OH)_4_), Wind, O_2_, MLD and log(Chl-a) with negative correlations (Figure 4e). Then, the environmental variables that covary most strongly with PC2 are: SSS, N*, O_2_, Wind and MLD on the positive side, *versus* log(Chl-a), log(Si(OH_4_)), and Si* on the negative side (Figure 4f). Therefore, the PCA indicates that the global gradient of temperature and productivity-related variables constitutes the first-order covariate of the PFGs’ species diversity. Meanwhile, the second-order covariate encapsulates gradients in nutrient ratios, oxygen concentration at depth, phytoplankton biomass and water mixing that separate oligotrophic gyres from upwelling systems, western boundary currents and minimum oxygen zones (Figure 4b,f). This second-order gradient covaries with the men annual richness of Coccolithophores, Jellyfish, Euphausiids, Appendicularians, Diatoms and Amphipods the most (Figure 4d).

The PCA also allowed us to investigate emergent Biodiversity-Ecosystem Functioning relationships as proxies of marine ecosystem services were also included as covariates. These proxies covary with the two PCs with varying strengths. Annually integrated megafauna diversity and mean annual net primary production (NPP) were positively correlated to PC1, whereas the e ratio (i.e., efficiency of the biological carbon pump) and the plankton size index (PSI) were negatively correlated with PC1. The mean annual amount of POC exported below the euphotic zone (FPOC_ex_), the PSI, chlorophyll-a concentration and NPP were the proxies scoring PC2. Therefore, regions in blue on Figure 4b indicate regions where the plankton communities are associated with productive conditions, larger phytoplankton cells and high POC fluxes at depth.

### 3.4 Uncertainties

Global mean uncertainty levels (Figure S3) showed significant variations across PFGs (Kruskal-Wallis rank sum test, Chi^2^ = 156454, p < 0.001). Most pairwise tests of uncertainty variations between PFGs (posthoc Dunn’s tests with a Bonferroni method for p-value corrections) showed significant variations (p < 0.001), except the following pairs (all p > 0.01): Coccolithophores x Poecilostomatoids, Jellyfish x Oithonids, Salps x Jellyfish, and Salps x Oithonids. Two PFGs have clearly higher levels of mean uncertainties in global richness estimates (Figure 5): Foraminifera (0.09 ± 0.06) and Appendicularians (0.08 ± 0.04). On the opposite end, Dinoflagellates, Diatoms and Calanoids show much lower mean uncertainty levels (all median uncertainties < 0.03). A strong negative linear relationship was found between uncertainty levels and sampling effort (Figure 5) indicating that uncertainty levels in mean annual richness decrease as more observations are available for fitting the SDMs. The mean uncertainty levels of Foraminifera and Appendicularians are above the standard error interval of the linear regression, suggesting the uncertainties of their richness estimates were higher than what can be predicted by sampling effort alone. On the opposite, Chaetognaths, Euphausiids and hyperiid Amphipods showed uncertainty levels lower than what can be predicted from sampling effort. Average uncertainty also decreased with the number of species modelled per group, but the linear fit was poorer than when considering sampling effort (adjusted R^2^ = 0.33, intercept = 5.75x10^-2^, slope = -1.93x10^-4^; p = 0.02).

**Figure 5:**
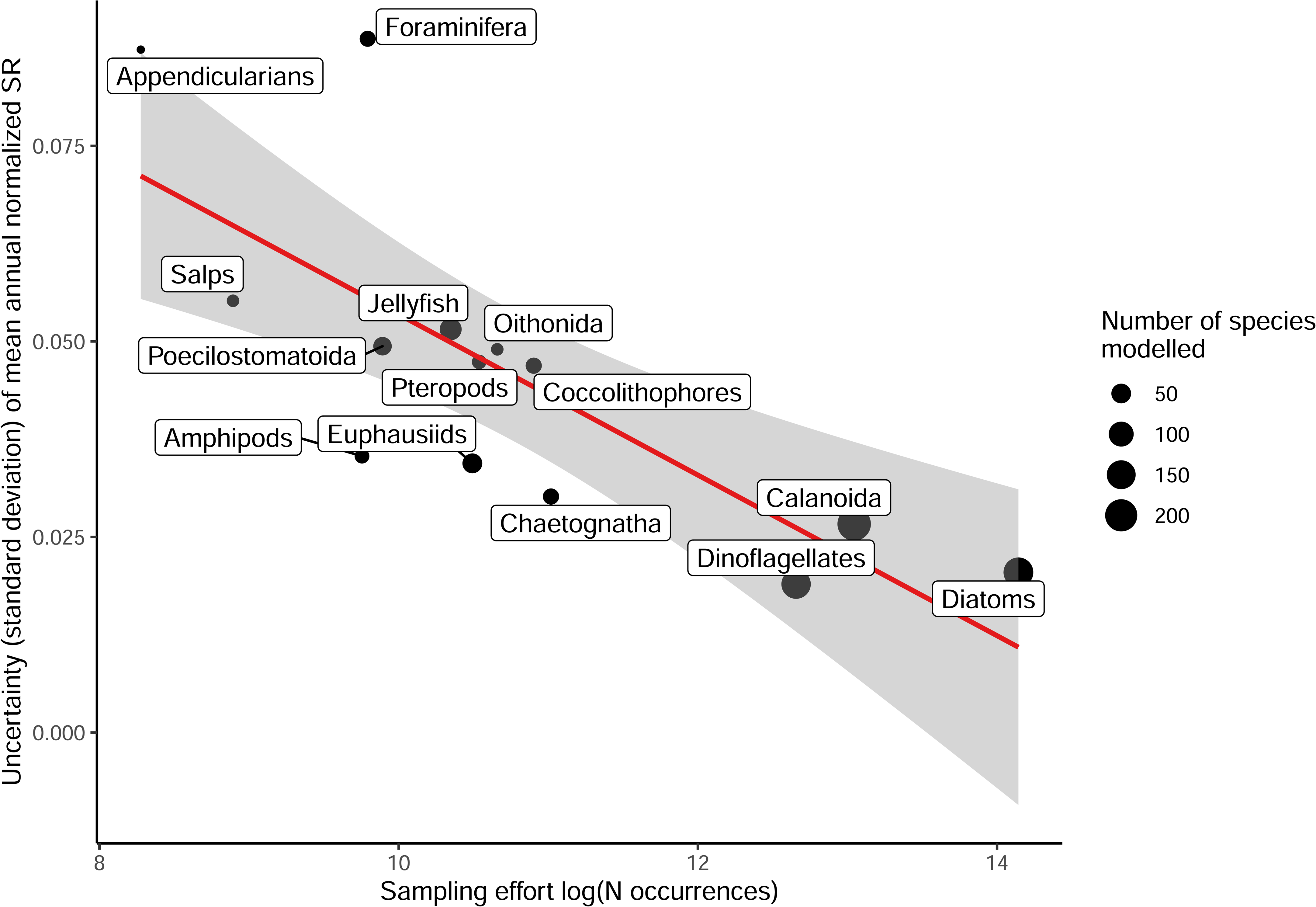
Negative linear relationship between sampling effort expressed as the logged number of species-level occurrences per plankton functional group (PFG) and the median uncertainty of the ensemble estimates of mean annual species richness (expressed in % of species modelled for each group). Global scale species richness uncertainty was estimated as the standard deviation of the mean annual richness across ensemble model members (n = 30). The values shown here correspond to the median of the PFG-level uncertainty for the global ocean. The size of the points shows the number of species modelled per PFG. The semi-transparent grey ribbon illustrates the standard error interval of the linear regression (n = 14, adjusted R^2^ = 0.55, intercept = 0.15, slope = -0.01; p-value = 0.001).

## 4. Discussion

### 4.1. Patterns and drivers of latitudinal diversity gradients across plankton functional groups

With their increasing species richness from the high latitudes to the low latitudes (Figures 2 & 3), all 14 modeled PFGs show latitudinal diversity gradients that are typical of marine ectotherms (Tittensor et al., 2010; Beaugrand et al., 2015). Perhaps even more notable is that none of the investigated groups shows a reverse gradient (Figure 2). This suggests that the underlying processes that control species diversity are common to widely different functional groups of plankton. Although all PFGs show decreasing diversity from the low to the high latitudes, they also show marked contrasts in the way the and the strength of the rate at which species accumulate with latitude (Figure 2). Pteropods, Poecilostomatoids, Oithonids and Salps show the steepest latitudinal diversity gradients while Amphipods, Appendicularians and the three phytoplankton groups show the weakest ones. The second component of the PCA (Figure 4b,d,f) captures the richness patterns that modulate those differences in latitudinal gradients (Figure 4a,c,e), permitting us to better analyze the covariance of PFG species richness with productivity-related variables (i.e., N*, Si*, chlorophyll-a, NPP; Figure 4f).

Among phytoplankton groups, the species diversity of Diatoms and Dinoflagellates show a stronger covariance with latitude than Coccolithophores diversity (Figure 2). Plus, the diversity of the latter decreases over a narrower latitudinal range (Figure 2; Document S1). Coccolithophores diversity is lower in eastern boundary upwelling systems and peaks in oligotrophic gyres (Figure 3c). Meanwhile, the latter emerge as more favorable regions for Diatoms and Dinoflagellates diversity, whose species richness dips in the high latitudes contrary to Coccolithophores (Figure 4b,d). This distinction stems from the way the SDMs captured the relationships between the occurrence data and the environmental predictors: Coccolithophores are the only PFG whose top ranking predictor was N* instead of SST (Figure 1), so the emergent Coccolithophore diversity patterns were less influenced by the latitudinal SST gradient compared to other PFGs (Figures 3 & 4). As a result, the strongest environmental covariates of Diatoms and Dinoflagellates diversity differ from those of Coccolithophores diversity (Figure 4f). Coccolithophores richness is positively associated with N*, i.e, the excess of nitrate over phosphate, and negatively associated with Si*, i.e., the excess of silicic acid over nitrate. The silicifying Diatoms show the opposite pattern as they are known to benefit from high silicic acid concentrations (Sarthou et al., 2005, Endo et al., 2018). We interpret this difference between Coccolithophores and Diatoms/Dinoflagellates as an emergent pattern stemming from their distinct ecological strategies and physiological traits. Most Coccolithophores (except species like *Emiliania huxleyi*; Paasche, 2001) are considered K-strategists characterized by higher nutrient affinity and lower growth rates that enable them to prevail in stable and oligotrophic conditions. Conversely, Diatoms and Dinoflagellates are usually considered as r-strategists that thrive in nutrient-replete and turbulent conditions thanks to their higher growth rates (Margalef, 1978). Such contrasting diversity patterns may emerge on a macroecological scale if Diatoms species recurrently outcompete Coccolithophores species under conditions of silicic acid replenishment, while being outcompeted under the conditions that lead to non-Redfield ratio uptakes of nitrogen (and higher N* values; Martiny et al., 2013). Therefore, the imprint of the distinct traits and ecological strategies on the biogeography (meaning the relationship between the species occurrences and the environmental predictors) of Coccolithophores and Diatoms/Dinoflagellates may be so strong that it was captured by our modelling approach. In situ observations (Smith et al., 2017; Endo et al., 2018) and ecosystem modelling studies (Nissen et al., 2018) support such resulting strong zonation of these two phytoplankton functional groups.

Among zooplankton groups, inter-groups differences along PC2 are less marked (Figure 4f) as most of these PFGs show similar latitudinal gradients in richness that mainly covary with temperature-related factors (Figure 4c,e). The strongest differences in mean annual richness are found between Jellyfish and Euphausiids versus Appendicularians and Amphipods. The former group has its highest species diversity in the subtropical gyres, while their richness dips in eastern upwelling systems and areas characterized by the presence of oxygen minimum zones (OMZs) at depth. In contrast, the species richness of the Appendicularians and Amphipods peaks in eastern upwelling systems and OMZs Figure 3; Document S2).

Amphipoda species richness also sits on the negative side of PC2 because it is not minimal in the Southern Ocean (Figures 2, 3 & 4). Most of the Amphipods modelled here belong to genera that mainly occur in high latitude ecosystems such as the Southern Ocean or the Arctic (e.g. *Themisto* spp.; Havermans et al., 2019). Therefore, we are confident that the relatively high mean annual richness of Amphipods at high latitudes reflects the imprint of the environment on their biogeography, likely due to the environmental filtering of their peculiar traits (but see Havermans et al., 2019). Most of the Amphipods modelled are known commensals and parasitoids of gelatinous plankton such as Jellyfish (Burridge et al., 2017), so their non-overlapping richness peaks along PC2 may seem surprising. It should be noted that most of the occurrences used here come from traditional plankton sampling techniques that are known for breaking the soft bodies of gelatinous zooplankton such as medusae and ctenophores. Therefore, our approach is unlikely to capture the imprint of fine scale parasitic relationships between gelatinous zooplankton and Amphipods on their species distribution and thus their large-scale species richness patterns.

The Appendicularians stand out because their species richness peaks in areas characterized by OMZs (Figure 3m; Document S2), a pattern that is hard to interpret because Appendicularians are not known to be particularly affiliated to such conditions. The recent global study by Soviadan et al. (2022) found pelagic tunicates (i.e. Salps and Appendicularians) to be more tolerant to the presence of OMZs than other zooplankton, in line with previous observations (Thuesen et al., 2005; Trueblood, 2019). We find oxygen to be an important predictor for Salps distribution (Figure 1) and we find Salps diversity to be weakly affected by the presence of OMZs contrary to all other zooplankton groups (Document 2), which could be a reflection of their relatively high tolerance to low oxygen conditions. Meanwhile, other PFGs that are known to be hypoxia-tolerant, like Jellyfish, Foraminifera, Oithonids and Poecilostomatoids (Thuesen et al., 2005; Haus et al., 2019; Soviadan et al., 2022) show lower richness in OMZs (Document S2). We put forward two hypotheses to explain such discrepancy: (i) method limitation, i.e., our approach may be limited in how well it can capture the imprint of low oxygen tolerances on the group-level biogeographies between Salps and other groups (because of group-level differences in data coverage for instance); (ii) species-level differentiation, i.e., the emergent patterns are driven by species-level differences in oxygen tolerances within the groups themselves (e.g. not all Jellyfish species are equally tolerant to low oxygen concentrations), which may lead to decreases in species-level habitat suitability indices, and thus a decrease in richness in areas where OMZs occur.

The drivers of plankton community assembly and diversity are debated as various studies identified different drivers, depending on whether they relied on traditional observations, metagenomics, or theoretical models (Dutkiewicz et al., 2020; Richter et al., 2021; Sommeria-Klein et al., 2021; Benedetti et al., 2021). Although our statistical approach does not allow to pinpoint the ecological mechanisms that determine the inferred latitudinal diversity gradients, the species-level ranks of relative importance (Figure 1) and the covariance analysis (Figure 4; Figure S4) support the view that temperature and the environmental factors covarying with it are the first-order controls on the distribution of species richness of the PFGs on a global scale. Therefore, on hand, our findings lend support to the kinetic energy hypothesis and the metabolic theory of ecology, according to which higher temperatures promote higher species diversity through increased speciation rates and the selection of warm-water tolerant species (Tittensor et al., 2010; Brown, 2014; Benedetti et al., 2021). On the other hand, trait-based ecosystem models found that various dimensions of global phytoplankton diversity are rather controlled by the rates of resource supplies, the grazing-mediated selection of functional types, and the turbulence associated with mesoscale and sub-mesoscale activity (Barton et al., 2010; Dutkiewicz et al., 2020). Other observational studies based on high-throughput DNA sequencing found local abiotic and biotic (e.g., biological interactions) factors and current-mediated dispersal to explain latitudinal patterns in plankton community structure (Richter et al., 2021; Sommeria-Klein et al., 2021). Here, environmental predictors related to climatic variability and turbulence at the scale of our study display either no correlations (e.g. annual SST range and EKE) or negative ones (e.g. MLD and Wind speed) with the PFGs mean annual richness (Figure 4; Figure S4). In addition, MLD, Wind speed and EKE do not surpass SST in terms of predictors’ rankings (Figure 1). The overall lack of positive relationships between mixing-related factors and species richness does not fall in line with the model-based studies that suggest that mixing and mesoscale activity promote the co-existence and evenness of PFGs (Barton et al., 2010; Dutkiewicz et al., 2020; Mangolte et al., 2022). These studies were conducted at a eddy-permitting (or eddy-resolving; Mangolte et al., 2022) resolution and incorporate competition processes between size-based PFGs, but are rarely evaluated against in situ data. Meanwhile, our approach is based on observations aggregated throughout decades to constrain empirical SDMs on spatial and temporal scales (1°x1°, monthly) that do not allow to resolve such competition effects, nor the impact of mesoscale features.

As already shown in terrestrial ecology, the imprint of various biological, ecological and climatic processes on species diversity is strongly scale dependent (Thuiller et al., 2015; Gonzalez et al., 2020). Therefore, we suggest that the discrepancies highlighted above are related to the scale-dependency of the various processes that structure marine plankton biodiversity. Different abiotic and biotic factors may emerge as dominant drivers of marine plankton species richness depending on the spatial and temporal scales covered by the data at hand.

### 4.2. Emergent Biodiversity-Ecosystem Functioning relationships

We find that the species richness of PFGs covaries positively with the species richness of oceanic fishes, sharks and mammals (Figure 4e,f), which implies that the diversity of PFGs and higher trophic levels are shaped by similar temperature-related drivers (Tittensor et al., 2010). This further supports the realism of our richness estimates as our SDMs ensemble seems to capture the main processes that constrain marine biodiversity on a global scale. Yet, we cannot here establish how the diversity of the smaller plankton influences the diversity of higher trophic levels. Second, we find that the efficiency of the biological carbon pump (i.e. the e-ratio) decreases with the species richness of all PFGs (Figure 4g,h). This finding is in line with the view that species-rich and functionally diverse communities are better able to retain the biomass produced within surface layers, whereas species-poor communities tend to leak higher rates of biomass out of the surface ocean. In addition, the PSI shows very similar correlation patterns to the e-ratio which implies that the emergent negative relationship between plankton species diversity and the efficiency of the biological carbon pump could also be mediated by changes in size structure. Indeed, the species-rich communities from the tropics tend to be composed of smaller species compared to the species-poor communities from the high latitudes (Uitz et al., 2010; Acevedo-Trejos et al., 2015, 2018; Brandão et al., 2021). High latitude communities tend to have higher proportions of large phytoplankton cells and large-bodied zooplankton whose functional traits are known to favor faster export of POC to depth (Stamieszkin et al., 2015; Tréguer et al., 2018; Henson et al., 2019; Brun et al., 2019). However, here, the absolute quantity of POC exported below the euphotic zone is covarying with PC2 and not PC1, meaning that it does not show strong correlation with the global species diversity gradients of most PFGs. It could be that the diversity of zooplankton groups does not strongly control the total amount of POC export but rather the fraction of NPP that is turned into export fluxes (Henson et al., 2019). More research is needed to help disentangle the effects of species richness from those of size distribution (or other size-related traits; Litchman et al., 2013) on export production and its efficiency.

The relationship between primary production and zooplankton diversity is unclear. Few zooplankton groups show strong loadings on PC2 which captures the global productivity gradient (Figure 4) and because log(Chl-a) ranks amongst the least important predictors of zooplankton species ranges (Figure 1; Table S3), meaning it was not included as a predictor in most of the SDMs. Although phytoplankton biomass is a determining factor for zooplankton biomass (Strömberg et al, 2009; Drago et al., 2022; Knecht et al., 2023), it could be less relevant for determining the spatial ranges of zooplankton species, and thus emergent species richness, if those are mainly controlled by temperature (Figure 1; Benedetti et al., 2021). We further examine the potential link with productivity by testing whether the PFGs show higher or lower mean annual richness within eastern boundary upwellings (EBUS), which represent relatively warm and productive areas (Chavez & Messié, 2009), compared to other tropical and less productive areas (Document S2). We find that most PFGs (except Salps, Amphipods and Poecilostomatoids) show significantly lower species richness in EBUS. This is interesting as it implies that conditions of higher productivity are not favorable to all species within PFGs, assuming that most zooplankton groups rely on primary production either directly (i.e. through grazing or filter feeding of particles) or indirectly (i.e. through the predation smaller grazing zooplankton). This finding supports the view that such conditions lead to environmental and biotic filtering (i.e. competitive exclusion) that select a subset of taxa at the expense of others, likely because of fitter traits combinations (Falkowski & Oliver, 2007; Litchman et al., 2010). Conversely, it contradicts the view that niche partitioning promotes the co-existence of various species within PFGs. As all three phytoplankton groups show lower species richness in EBUS (Document S2) and in productive temperate areas characterized by seasonal bloom regimes (Figure 4; Righetti et al., 2019), our results suggest that, at the scale of our study, higher productivity might be carried by subsets of the phytoplankton community.

Ultimately, the richness of phytoplankton groups decreases with higher nutrient availability which can be interpreted through two mutually non-exclusive processes. On one hand, species-rich communities draw down nutrient concentrations more efficiently, meaning that phytoplankton diversity optimizes nutrients use efficiency in the system (Falkowski & Oliver, 2007; Marañón, 2015). This would translate into lower relative rates of biomass exported from surface ecosystems, which is corroborated by the emergent negative correlations between phytoplankton species richness and the e ratio (Figure 4). On the other hand, systems such as tropical gyres that are characterized by weaker seasonal variations and lower nutrient availability might sustain species-rich communities through niche partitioning that enables the co-existence of many taxa characterized by different functional traits (Litchman et al., 2010; Edwards et al., 2016). Yet again, we underline that the scale of our study may not align with the finer spatio-temporal scales at which such processes occur, thus masking their influence. Under either hypotheses, our results point towards the existence of links between key ecosystem functions (i.e. nutrients use efficiency and carbon export) and the species diversity of PFGs. Our results warrant further field studies examining the relationships between ecosystem functions, species richness and community traits expression, as well as the verification and extension of existing ecological theory (Chust et al. 2018).

### 4.3. Uncertainty sources and caveats

In this study, PFGs had to be defined based on our current knowledge of the ecological and biogeochemical functions associated with each broad taxonomic group, as an exhaustive compilation of functional trait across multiple plankton species is still missing. One of the main caveats of this approach is that we are not accounting for species-level and organism-level variations in functional traits within PFGs. Within groups such as Diatoms, Jellyfish or Salps, species’ sizes span one or several orders of magnitude (Lucas et al., 2011; Henschke et al., 2016; Leblanc et al., 2018). Since most functional traits scale with size (Litchman et al., 2013), our approach underestimates the existence of the smaller functional groups nested within those chosen here (Leblanc et al., 2018; Benedetti et al, 2022). Recent functional trait syntheses enabled to identify at least eleven functional groups of copepods nested within the three groups used here (Benedetti et al., 2022). Such studies remain impossible to perform across more than ten PFGs as long as we are lacking a “panplankton” synthesis of functional traits for phyto- and zooplankton groups (Barton et al., 2013; Litchman et al, 2013; Martini et al., 2021). This caveat is more relevant for those PFGs characterized by large species richness (n > 50; i.e. Dinoflagellates, Diatoms, Calanoids, Jellyfish and Euphausiids) as smaller PFGs are likely to display a much lower variability in traits. Our definition of PFGs aims to achieve an optimal trade-off between established inter-groups differences in size classes and ecological or biogeochemical functions (Le Quéré et al., 2005; Hood et al., 2006; Heneghan et al., 2020). We are confident that the present PFGs capture distinct functional entities, at least from the inter-group point of view.

The other caveats of our approach are those that are inherent to the use of SDMs (Elith & Leathwick, 2009; Thuiller et al., 2019). SDMs do not incorporate some of the processes that shape plankton distributions, such as dispersal, biotic interactions and population-level variations in niches (Elith & Leathwick, 2009). The present SDMs assume that plankton species distributions can be reproduced as a function of the combinations of environmental predictors defining a species’ niche. Such assumptions remain supported on macroecological scales where the imprint of biological interactions is considered to be relatively small (Araújo & Rozenfeld, 2014; Thuiller et al., 2015), and where connectivity through sea currents is high enough to support weak dispersal limitations (Jönsson & Watson, 2016). There is now mounting evidence that the spatial ranges of marine ectotherms are mainly shaped by their thermal niches (Pinsky et al., 2019; Fredston et al., 2021) and that empirical distribution models robustly reproduce the distributions of marine ectotherms (Sunday et al., 2012). As our ensemble of SDMs can reproduce the latitudinal diversity gradients observed for several groups, we are confident in the present estimates of mean annual richness (Benedetti et al., 2021; Righetti et al., 2023).

Nonetheless, we carefully investigated the level uncertainties associated to the latter (Figure 5; Figure S3). Previous work showed that SDMs choice drives the variability between model projections (Benedetti et al., 2017; Thuiller et al., 2019). We show that there are strong differences in uncertainty levels between PFGs, with Foraminifera and Appendicularia showing the most uncertain species richness estimates, and Diatoms, Dinoflagellates and Calanoids showing the least uncertain (Figure 5). Consequently, we believe that the present richness patterns should be taken with caution for Foraminifera and Appendicularia. PFG-level variations in species richness uncertainty could be robustly explained by differences in data coverage and species richness (Figure 5). We acknowledge that these two factors are not independent since richer PFGs tend to have a larger pool of occurrences available for model training (Figure S2). The diversity estimates of Foraminifera and Appendicularia (Figures 2 & 3) can thus be largely improved through our approach by enriching the spatial coverage of their occurrence pool which is currently relatively low. A wealth of foraminifera species occurrences derived from sediment records is available (Siccha & Kucera, 2017), but they could not be included in our study as we focus on samples taken in the upper water column to be consistent across all PFGs. Several studies used such sediment records to estimate pre-industrial and modern planktic Foraminifera diversity gradients (Lombard et al., 2011; Jonkers et al., 2019; Yasuhara et al., 2020). They usually found Foraminifera diversity to peak northwards of the subtropics (∼ 40° latitude). Therefore, the fact the we find Foraminifera richness to peak between 0° and 20° latitude could be driven by limited data availability.

We underline that we aimed to model species richness gradients and not absolute species richness values since many species could not be modelled due to limited occurrence numbers. Here, we managed to model about 20% of the 1704 species recorded in PhytoBase (Righetti et al., 2020) and 15% of the 3206 species recorded in ZooBase (Benedetti et al., 2021). As a consequence, some of the present richness patterns are likely underestimated in regions of the ocean where lower sampling effort may have led to incomplete sampling of the plankton community (Figure S2). The species modeled here are those that are the most frequently detected by traditional sampling and identification techniques either because they dominate the community in terms of abundance and biomass, or because they correspond to relatively large species that are more easily sampled and identified by taxonomists (Table S1). Ser-Giacomi et al. (2018) showed that the ranges of rare and non-dominant plankton species exhibit no biogeographical signature, meaning that they are hardly relatable to environmental gradients. Consequently, there is no guarantee that such rare taxa can be robustly modelled through our numerical approach. Nevertheless, the recent study by Righetti et al. (2023) showed that the SDM framework used here is able to robustly reproduce observed global species richness gradients from scattered data. Finally, the present diversity patterns are based on species richness (Figures 2 & 3) and not an abundance-weighted diversity index (e.g, Shannon-Wiener index, like in Barton et al., 2010). Therefore, we are likely overestimating true diversity in areas where a subset of species dominate community abundance and biomass. The steepness of some of the present latitudinal gradients may be: (i) underestimated if diversity is overestimated towards high latitudes, or (ii) overestimated if diversity is overestimated in the tropics and near the equator.

## 5. Conclusions

Our study provides estimates of global ocean species richness and composition that were obtained in a coherent and comparable manner based on state-of-the-art distribution modelling across 14 PFGs. We find that all PFGs display global diversity gradients that are likely driven by processes related to the latitudinal temperature gradient. This leads to richer communities in the low latitudes compared to the high latitudes, similar to what is observed for higher trophic levels (Tittensor et al., 2021). Yet, individual PFGs show substantial differences in terms of the detailed pattern in mean annual species richness, with some groups having their maxima at the equator, and others near 30° latitude. We interpret this to be a consequence of inter-group or intra-group competition and/or selection effects that imprint the biogeography of the individual PFGs. We put forward the idea that the dominant processes underlying marine plankton diversity structure are scale-dependent, with temperature playing an important role on macroecological scales (e.g., planetary scale, basin scale, decadal scale), while other factors such as biotic interactions and dispersal may dominate on smaller scales (e.g., subregional to local scales, weekly scales).

Our study also supports the existence of links between ecosystem functions and the diversity of PFGs. We find that the efficiency of the biological carbon pump covaries negatively with plankton species richness, supporting the view that species-rich and functionally diverse communities are better able to recycle the biomass and nutrients within surface layers. Most PFGs show lower diversity in conditions of higher productivity, as those emerged as not favorable to all species within each PFG. Our study thus supports the view that and biotic filtering (i.e. competitive exclusion) takes place in productive conditions as a subset of taxa are selected likely because of fitter traits combinations.

We hope that our study will be used as a basis to bridge the gap between observational and model-based studies and to identify the scaling laws that describe which drivers of plankton diversity govern community assembly. The development of modern imaging and DNA sequencing techniques will allow to get more complete listings of plankton species and their associated traits (Sunagawa et al., 2015; de Vargas et al., 2015), yet data availability is still too limited to investigate the scale dependence of the drivers of diversity and community assembly. Understanding how these new data types can be integrated in the present modelling framework and into new macroecological theories will be determinant to build a unifying and holistic vision of marine plankton biodiversity and its effects on ecosystem functions (Cardinale et al., 2012; Gonzalez et al., 2020).

## Acknowledgements

We thank all contributors involved in the plankton species field sampling and identification and we acknowledge the efforts made to share such data through publicly available online archives.

## Data Archiving

All the plankton species occurrences underlying the main results of this paper are publicly available online. See: https://doi.pangaea.de/10.1594/PANGAEA.904397 for the phytoplankton data. See: https://zenodo.org/record/5101349#.YSUOjMY69pQ for the zooplankton data. All R codes used to generate the results of the study are stored on the GitHub account of F.B. (https://github.com/benfabio).

