## Supplemental Document 1 for "Global gradients in species richness of marine plankton functional groups"

**Document S1 (below): Zonal distribution of mean annual richness (expressed in % of species modelled) and its first derivative as a function of absolute latitude for the 14 plankton functional groups (PFGs).**

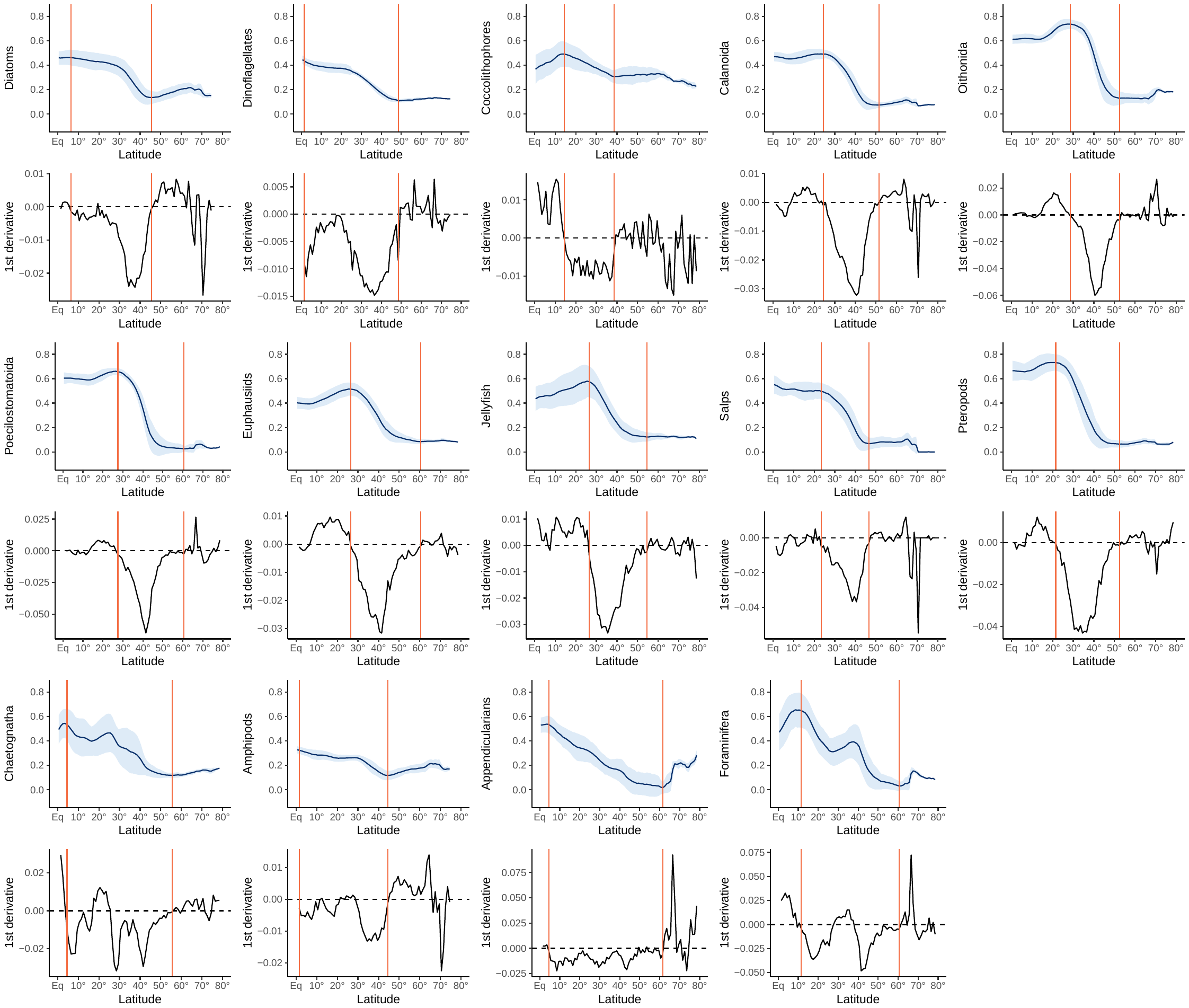

**Figure**: Zonal distribution of mean latitudinal richness (expressed in % of species modelled) and its first-order derivative as a function of absolute latitude (1° resolution) for the 14 plankton functional groups (PFGs) modelled. The blue curves illustrate the latitudinal averages of species richness per absolute degrees of latitude. The light blue ribbons illustrate the standard deviation associated with the latitudinal averages (i.e., longitudinal variability). The solid vertical orange lines indicate the range of absolute latitude of interest over which linear regressions were computed to estimate the rate of decrease in species accumulation from the tropics to the high latitudes.

The groups’ species richness estimates are normalized to the number of species modelled so they range between 0 and 1 with values equal to 1 indicating the grid cells where environmental conditions allow all species from the same PFG to be modelled as present (i.e., species accumulation). Based on these estimates, we explored the inter-PFGs differences in latitudinal diversity gradients by estimating the rate of the decrease in species accumulation from the most diverse regions (i.e., the tropics) to the least diverse ones (i.e., the sub-poles and the poles). We fitted linear regressions between mean latitudinal species richness and absolute latitude. The linear regressions were not computed across the full latitudinal range as most PFGs displayed non monotonic and nonlinear latitudinal gradients of richness (see Figure above). Here, we aimed to compare PFGs on the basis of their relative rate of species accumulation, rather than trying to prove the existence of linearity among the latitudinal gradients. Therefore, we computed the first derivatives of the mean latitudinal annual richness over absolute latitude to detect the latitudinal ranges over which the groups’ richness shows a monotonic and nearly linear decrease in species richness (Figure above). Latitudes > 65° were excluded since changes in diversity beyond this threshold only occur within a very small (often < 0.1) range of species richness. The linear regressions were fitted from the first absolute latitude that is followed by ten consecutive negative derivative values (i.e., a monotonic richness decrease) to the first latitude showing a positive derivative (i.e., richness increase). Two PFGs (Chaetognaths and Foraminifera) showed a local maximum (i.e., ten consecutive positive derivative values) between two phases of richness decrease. We chose to integrate these local maxima into the linear regressions instead of fitting two smaller and separate linear models for the same PFG.

The Table below summarizes the coefficients and the skill of each linear regression and the minimum and maximal latitudes over which they were computed for each PFG.

| PFG | Minimum latitude | Maximum latitude | Slope | Intercept | R^2^ |
| --- | --- | --- | --- | --- | --- |
| Diatoms | 6.5° | 45.5° | -0.0087 | 0.5819 | 0.8374 |
| Dinoflagellates | 1.5° | 48.5° | -0.0071 | 0.4794 | 0.9023 |
| Coccolithophores | 14.5° | 38.5° | -0.0079 | 0.6143 | 0.9972 |
| Calanoids | 24.5° | 51.5° | -0.0195 | 1.0126 | 0.9621 |
| Oithonids | 28.5° | 52.5° | -0.0321 | 1.7435 | 0.9493 |
| Poecilostomatoids | 27.5° | 60.5° | -0.0239 | 1.3171 | 0.8873 |
| Euphausiids | 26.5° | 54.5° | -0.0151 | 0.9110 | 0.9236 |
| Jellyfish | 26.5° | 54.5° | -0.0182 | 1.0213 | 0.9135 |
| Salps | 23.5° | 46.5° | -0.0217 | 1.0627 | 0.9135 |
| Pteropods | 21.5° | 52.5° | -0.0268 | 1.3423 | 0.9458 |
| Chaetognaths | 4.5° | 55.5° | -0.0082 | 0.5720 | 0.8859 |
| Amphipods | 1.5° | 44.5° | -0.0041 | 0.3393 | 0.8329 |
| Appendicularians | 4.5° | 61.5° | -0.0094 | 0.5377 | 0.9781 |
| Foraminifera | 11.5° | 60.5° | -0.0118 | 0.7177 | 0.8832 |

The slopes of the linear regressions were then used to compare the steepness of the latitudinal diversity gradient (i.e., rate of species accumulation over latitude) between PFGs through covariance analysis (ANCOVA). ANCOVA reveals that the coefficients of the linear regressions fitted vary significantly between PFGs (F = 1256.11; p < 0.001), indicating that most PFGs show different rates of decrease in species accumulation along the monotonic parts of the richness gradients. Posthoc Tukey’s multi-comparison tests indicate that the following pairs of PFGs show significant differences in the coefficients of the linear models (all p < 0.001): Oithonida – Diatoms; Poecilostomatoida – Diatoms; Euphausiids – Diatoms; Jellyfish – Diatoms; Pteropods – Diatoms; Amphipods – Diatoms; Coccolithophores – Dinoflagellates; Calanoida – Dinoflagellates; Oithonida – Dinoflagellates; Poecilostomatoida – Dinoflagellates; Euphausiids – Dinoflagellates; Jellyfish – Dinoflagellates; Salps – Dinoflagellates; Pteropods – Dinoflagellates; Chaetognatha – Dinoflagellates; Amphipods – Dinoflagellates; Foraminifera – Dinoflagellates; Oithonida – Coccolithophores; Amphipods – Coccolithophores; Appendicularians – Coccolithophores; Oithonida – Calanoida; Amphipods – Calanoida; Appendicularians – Calanoida; Poecilostomatoida – Oithonida; Euphausiids – Oithonida; Jellyfish – Oithonida; Salps – Oithonida; Pteropods – Oithonida; Chaetognatha - Oithonida; Amphipods – Oithonida; Appendicularians – Oithonida ; Foraminifera – Oithonida; Chaetognatha – Poecilostomatoida; Amphipods – Poecilostomatoida; Appendicularians – Poecilostomatoida; Chaetognatha – Euphausiids; Amphipods – Euphausiids; Appendicularians – Euphausiids; Chaetognatha – Jellyfish; Amphipods – Jellyfish; Appendicularians – Jellyfish; Amphipods – Salps ; Appendicularians – Salps ; Chaetognatha – Pteropods ; Amphipods – Pteropods ; Appendicularians – Pteropods ; Amphipods – Chaetognatha; Appendicularians – Chaetognatha ; Appendicularians – Amphipods ; Foraminifera – Amphipods ; Foraminifera – Appendicularians.
