## Supplemental Document 2 for "Global gradients in species richness of marine plankton functional groups"

**Document S2: How do OMZs and EBUS affect the SR of PFGs?**

Oxygen minimum zones (OMZ) and eastern boundary upwellings (EBUS) are major biogeochemical features of the ocean whose impacts on plankton biodiversity remain poorly known (Chavez & Messié, 2009; Wishner et al., 2018; 2020), especially when compared to the impact of the temperature gradient (Tittensor et al., 2010; Benedetti et al., 2021). Therefore we aim to assess the impact of these features on the species richness (SR) of the plankton functional groups (PFGs). To do so, we binarily classify (1/0) each grid cell of the ocean as an “OMZ cell” or an “EBUS cell” (Figure 1 below), and perform two-sided Wilcoxon tests to examine whether OMZ and EBUS cells display higher or lower levels of PFGs SR compared to other cells from comparable latitudinal ranges. For OMZ, only the zooplankton PFGs are considered as those are the groups whose SR is likely to be impacted by the presence of an underlying OMZ at depth.

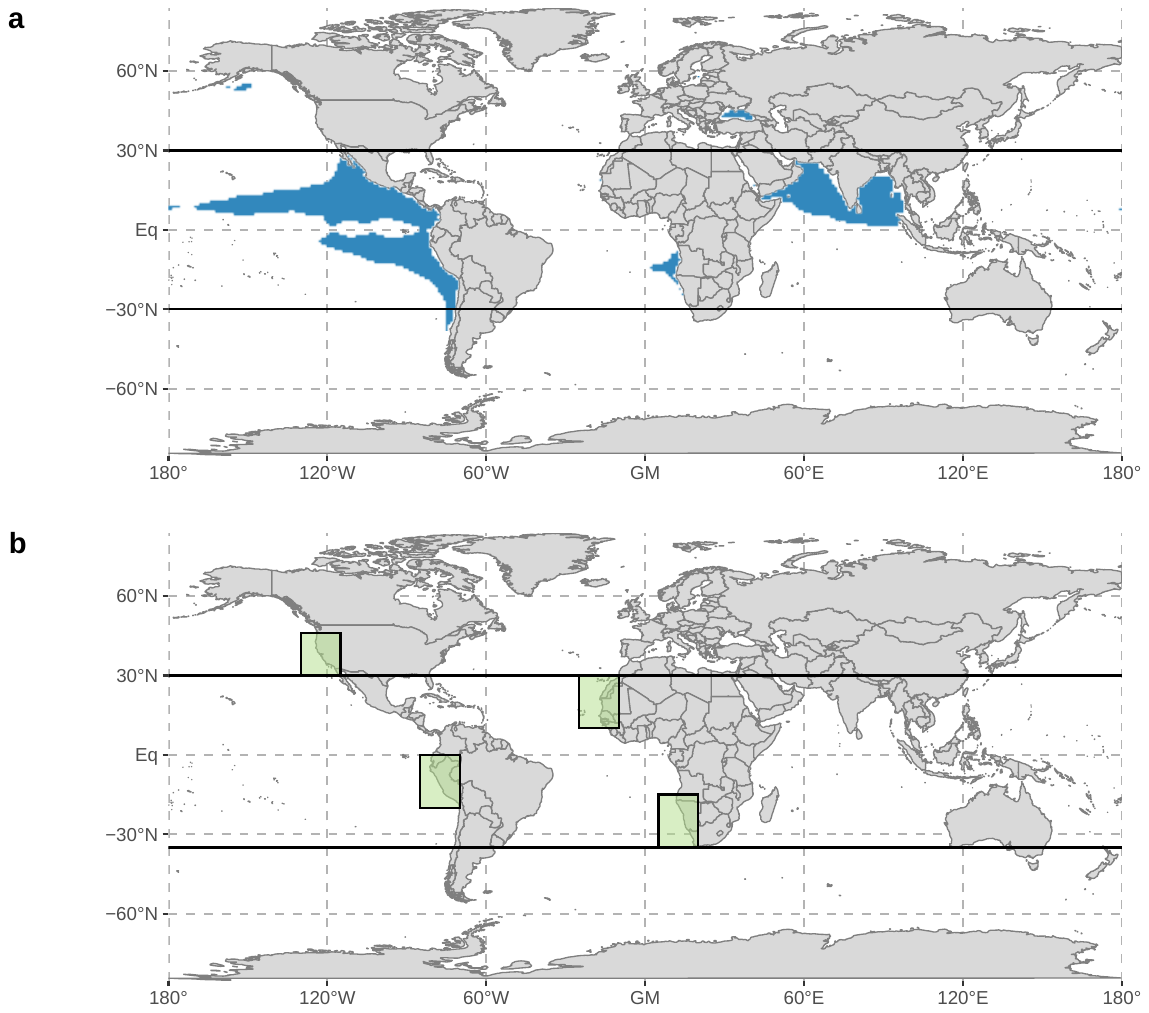

**Figure 1**: Geographical position of the a) oxygen minimum zones (OMZ) and b) eastern boundary upwellings (EBUS) defined to test their potential relationship with the mean annual species richness of the plankton functional groups (PFGs) modelled in our study. OMZ were defined based on the 61.0 µmol/kg at 200m depth criterion of Hofmann et al. (2011). The bounding boxes of the EBUS come from Chavez & Messié (2009).

OMZ cells correspond to those surface cells of the WOA exhibiting a mean annual dO_2_ value < 61.0 µmol/kg at 200m depth (Hofmann et al., 2011). Various depth thresholds (100, 200, 300, 400 and 500m) were explored to check the robustness of the tests results to this criterion. Here, we do not explicitly test how the presence of an OMZ at 200m affects zooplankton SR since our SR estimates are valid for the surface conditions of the open ocean. Rather, we test whether our modelled patterns of surface SR support the view that OMZs should lead to a decrease in zooplankton SR as only a subset of species characterized by traits enabling can withstand limited oxygen availability at depth (Wishner et al., 2018; 2020). For the Wilcoxon tests, only those non-OMZ cells falling in the tropical band (i.e., absolute latitude < 30°) are considered to: (i) balance sample sizes between OMZ (n = 2243) and non-OMZ cells (n = 12857) and (ii) remove the spatial variability in SR that is driven by the strong temperature gradient between the tropics and the poles, as this is known to be the main driver of marine latitudinal diversity gradients (Tittensor et al., 2010; Benedetti et al., 2021).

EBUS cells are those cells falling in the spatial boundary boxes defined by Chavez & Messié (2009) for the four main EBUS: Peru, California, North-West Africa and Benguela (Figure 1). As with OMZ, we cared to remove the variability in SR that would be mainly driven by SST to focus on the potential impact of EBUS conditions. Therefore, the California EBUS is removed from the analysis as it is much colder than the other three EBUS. In addition, only those non-EBUS cells whose latitude fall between the upper and lower latitudinal limits of the three EBUS considered (i.e., 34°S and 30°N) are considered for the variance analysis in order to balance sample sizes. Because non-EBUS cells (n = 15517) were still nearly 30 times more numerous than EBUS cells (n = 516), 999 Wilcoxon tests were performed by randomly subsampling 500 of both cell types to verify sample size differences did not bias our findings.

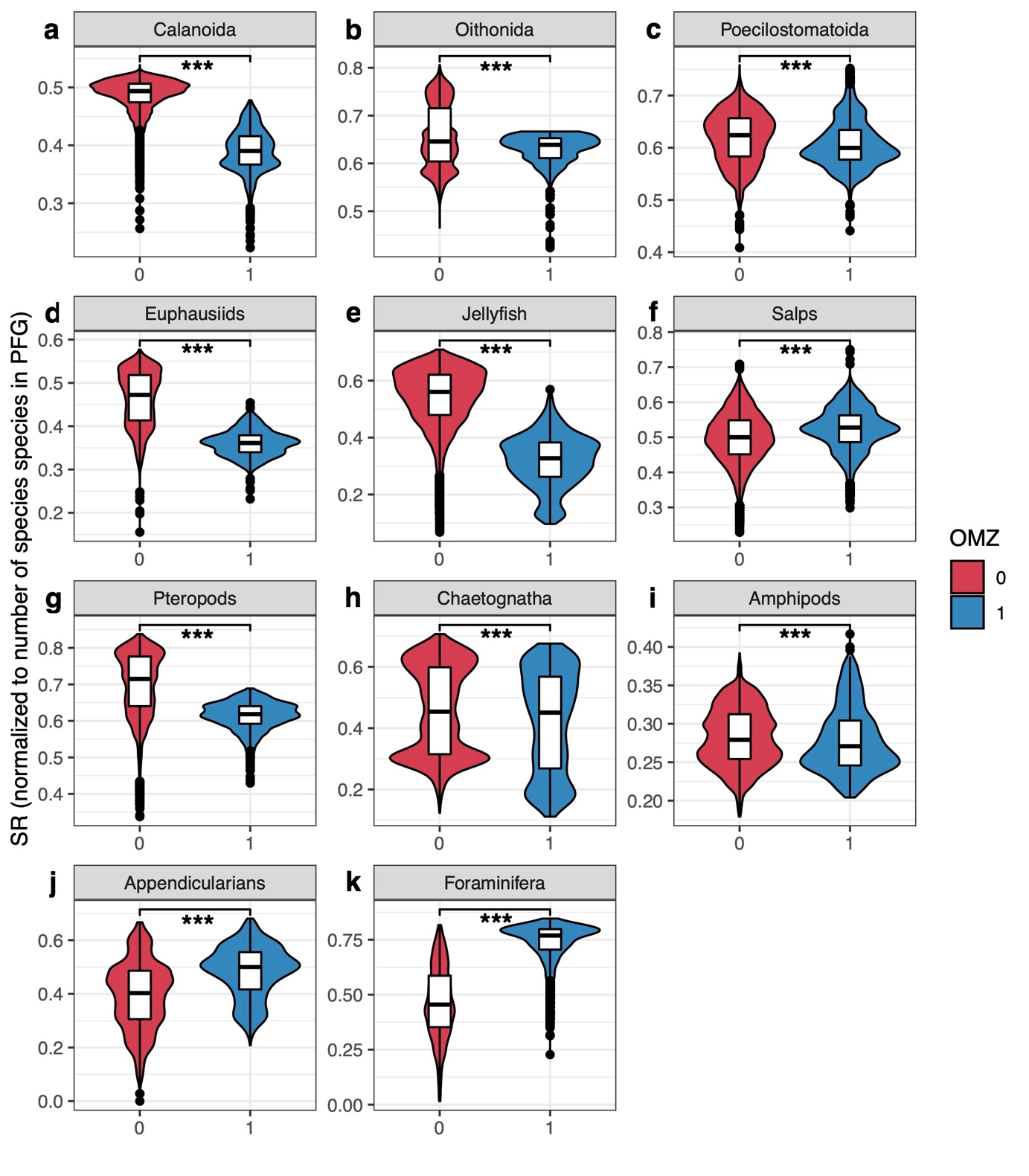

**Figure 2**: Distribution of mean annual species richness (SR, expressed in % of species modelled) between OMZ cells (blue, n = 2243) and non-OMZ cells (red, n = 12857) for a) Calanoida, b) Oithonida, c) Poecilostomatoida, d) Euphausiids, e) Jellyfish, f) Salps, g) Pteropods, h) Chaetognatha, i) Amphipods, j) Appendicularians, and k) Foraminifera. Only grid cells falling in the tropical band (i.e., latitude < 30°) were considered. The violin plots indicate grid cells density along the SR gradient. The central horizontal lines indicate median values, the boxes illustrate the interquartile ranges and the error bars indicate 5^th^ and 95^th^ percentiles. Differences in mean annual SR between OMZ and non-OMZ cells were tested through Wilcoxon tests (alpha = 0.01). All tests displayed p-values < 0.01 even when randomly re-sampling grid cells to balance sample sizes.

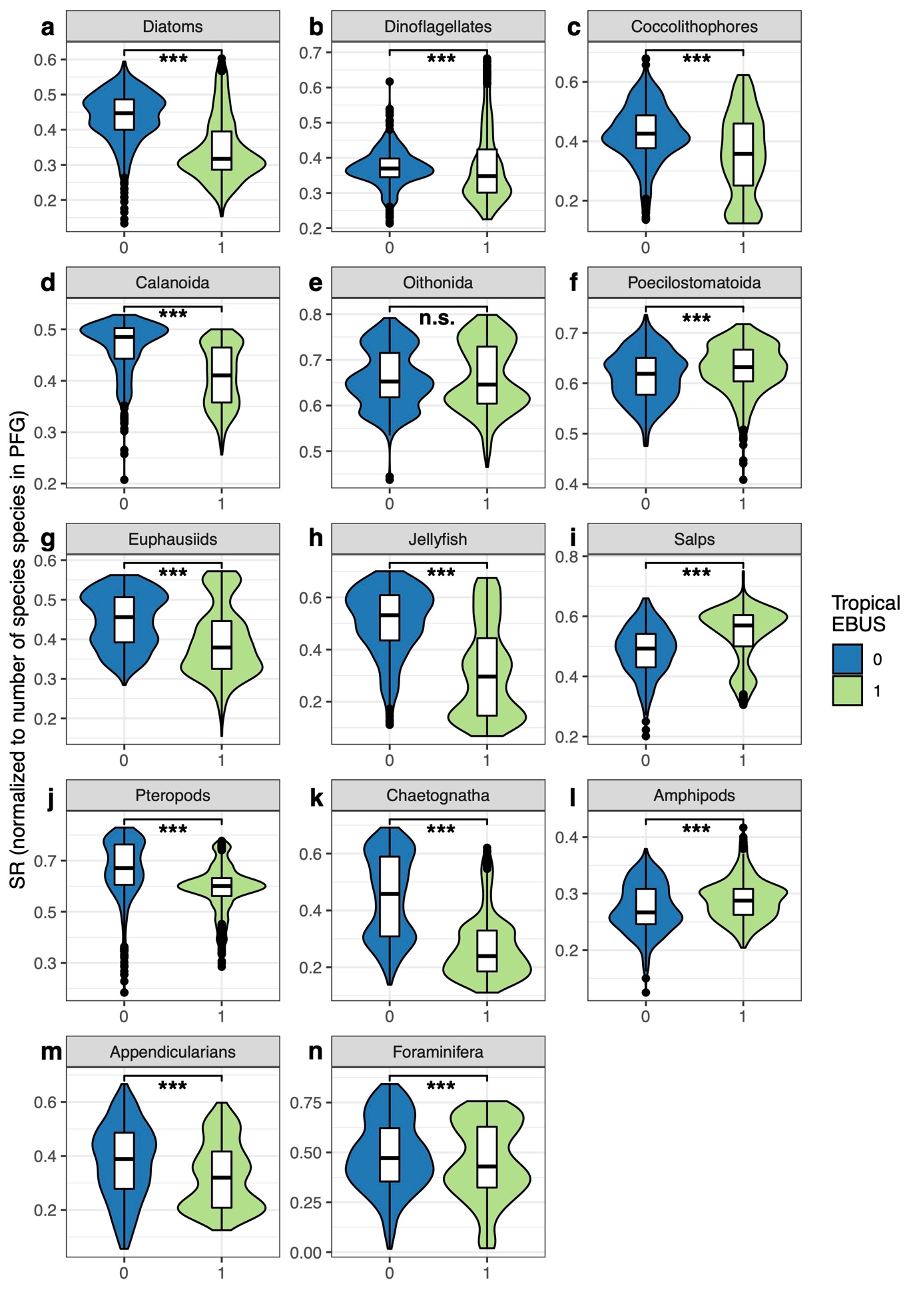

**Figure 3:** Distribution of mean annual species richness (SR, expressed in % of species modelled) between EBUS cells (green, n = 516) and non-EBUS cells (blue, n = 15517) for d) Calanoida, e) Oithonida, f) Poecilostomatoida, g) Euphausiids, h) Jellyfish, i) Salps, j) Pteropods, k) Chaetognatha, l) Amphipods, m) Appendicularians, and n) Foraminifera. Only grid cells falling in the tropical band (i.e., latitude < 30°) were considered. The violin plots indicate grid cells density along the SR gradient. The central horizontal lines indicate median values, the boxes illustrate the interquartile ranges and the error bars indicate 5^th^ and 95^th^ percentiles. Differences in mean annual SR between EBUS and non-EBUS cells were tested through Wilcoxon tests (alpha = 0.01). All tests displayed p-values < alpha, except Oithonida which always showed p-values > alpha, even when randomly resampling grid cells to balance sample sizes.

Nearly all zooplankton groups show a significant decrease in mean annual SR between tropical OMZs and non-OMZs cells except foraminifera, appendicularians and salps whose SR increased in OMZs cells (Figure 2 above). The strength of those SR variations differ between PFGs, as indicated by the difference in median SR between OMZs and non-OMZs cells given in brackets below. Jellyfish show the strongest decrease in median SR (-0.23), followed by euphausiids (-0.11), calanoid copepods (-0.10) and pteropods (-0.09). Meanwhile, foraminifera display increase in median SR (+0.31) in OMZs cells. The median SR of appendicularians and salps also increase but more moderately (+0.10 and +0.03, respectively). The remaining four PFGs show very low decreases in SR (all differences in median SR < -0.02).

Similarly, we test the impact of tropical EBUS on the mean annual SR of the 14 PFGs (Figure 3 above). All PFGs but oithonid copepods show significant variations in mean annual SR between tropical EBUS and non-EBUS cells. Salps, hyperiid amphipods and poecilostomatoid copepods display moderate increases in median SR (+0.08, +0.02 and +0.01, respectively). Again, jellyfish display the strongest decrease in median SR (-0.22), followed by chaetognaths (-0.20), diatoms (-0.13), appendicularians and pteropods (-0.08 median SR for both). The remaining five PFGs display very low decreases in SR (all differences in median SR < -0.07).
