## Supplemental Figure 1 for "Global gradients in species richness of marine plankton functional groups"

**Figure S1**: Relative contribution of each plankton functional group (PFG) to the total number of species modelled (n = 845).

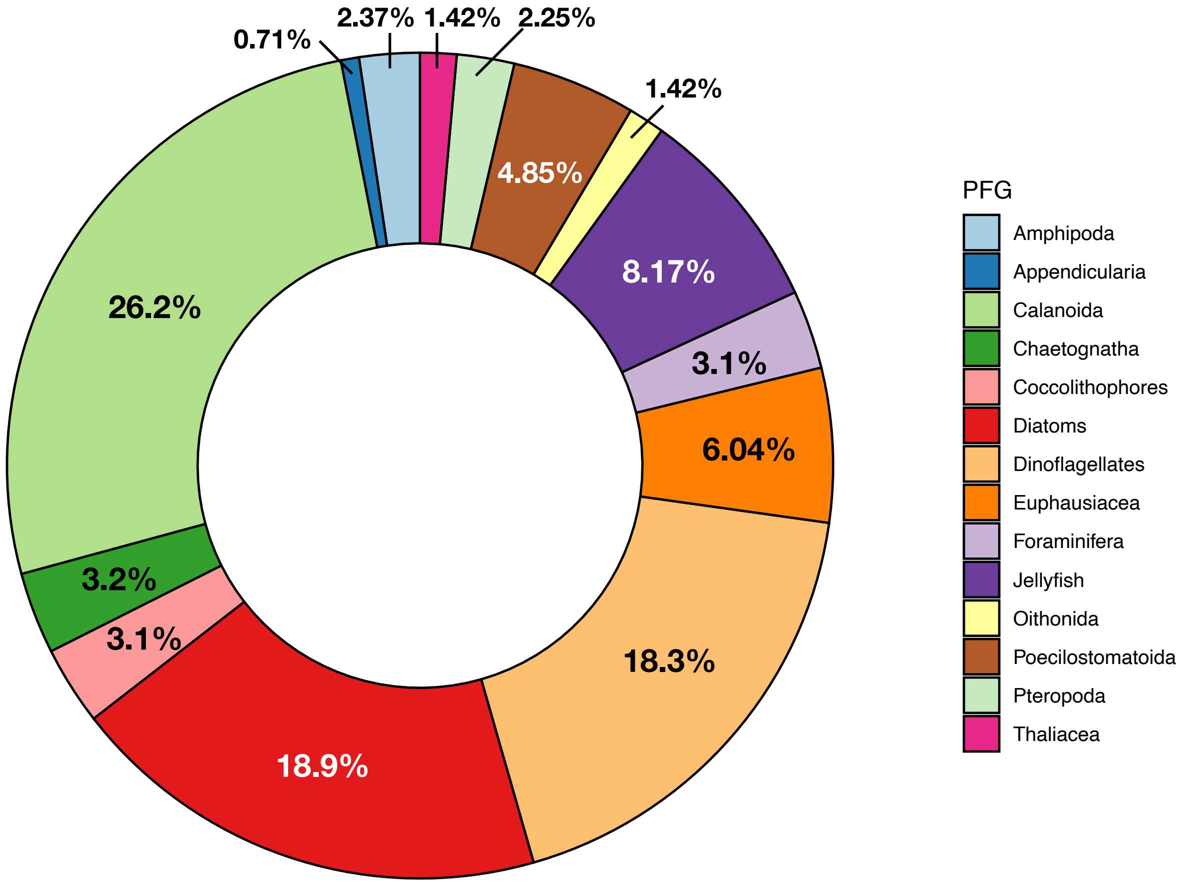
