## Supplemental Figure 2 for "Global gradients in species richness of marine plankton functional groups"

**Figure S2**: Spatial distribution of sampling effort (log of N unique species occurrences) for: a) Diatoms, b) Dinoflagellates, c) Coccolithophores, d) Calanoida, e) Oithonida, f) Poecilostomatoida, g) Euphausiids, h) Jellyfish, i) Thaliacea, j) Pteropods, k) Chaetognatha, l) Amphipods, m) Appendicularians, and n) Foraminifera.

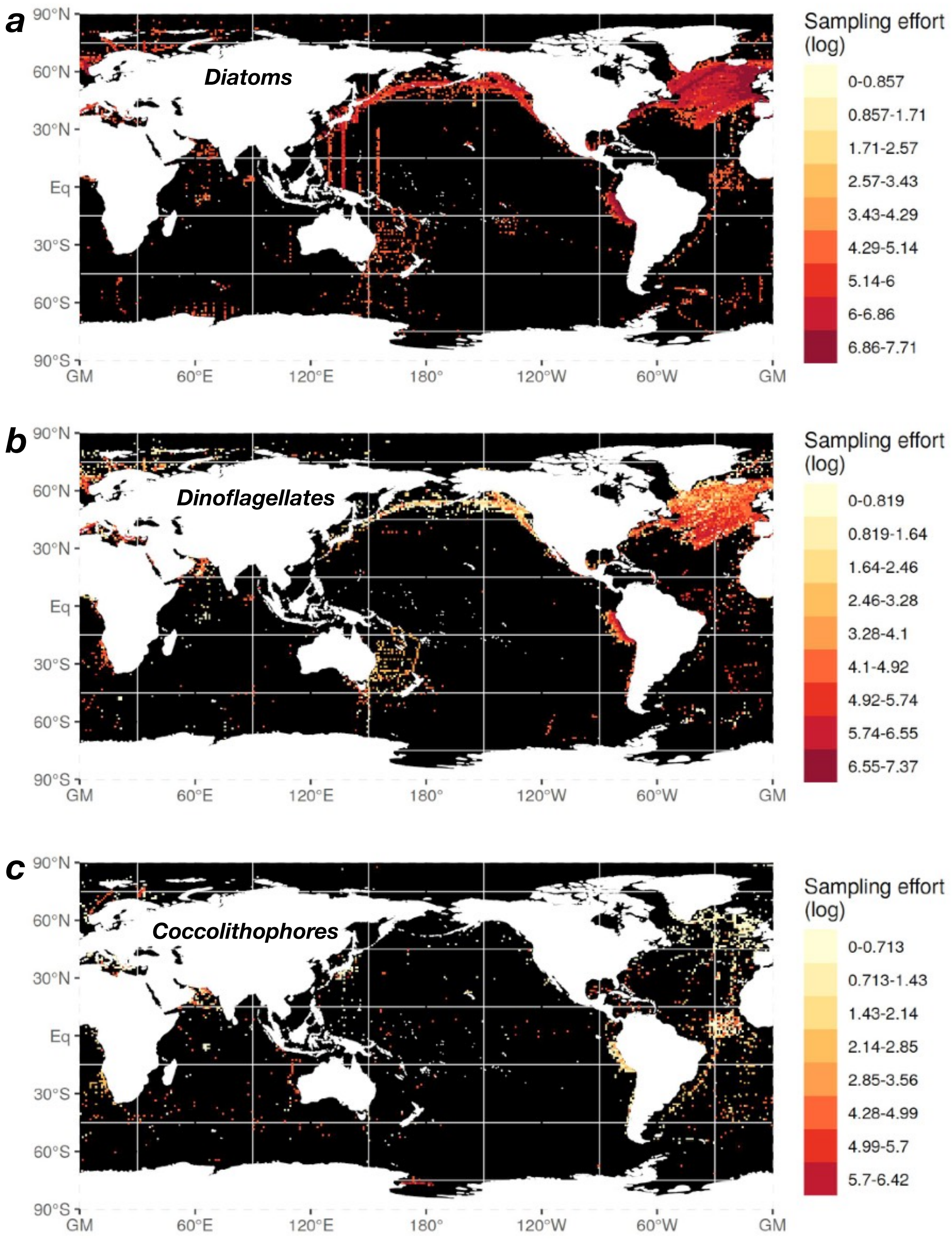

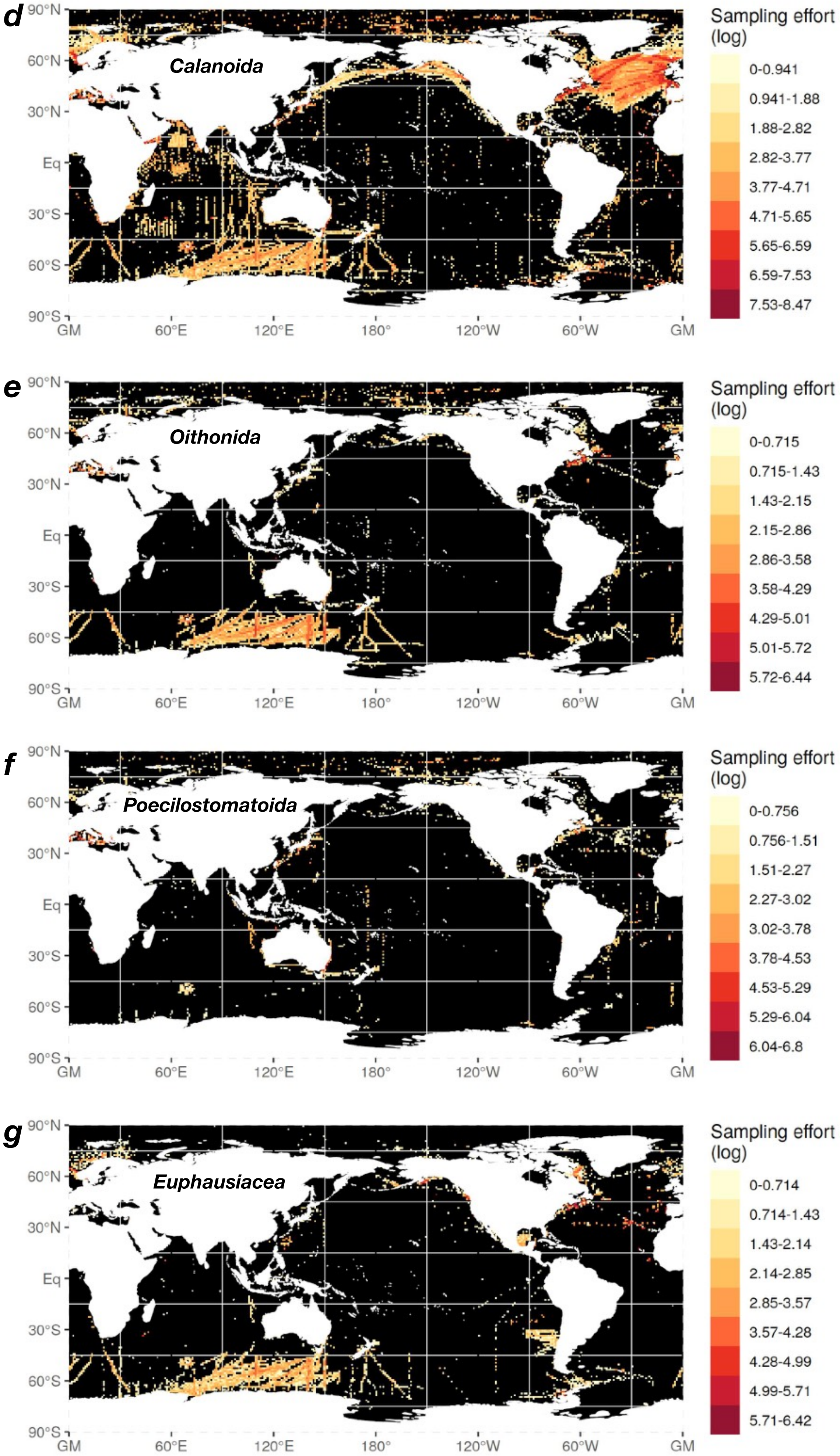
d
Calanoida
Sampling effort (log)
0-0.941
0.941-1.88
1.88-2.82
2.82-3.77
3.77-4.71
4.71-5.65
5.65-6.59
6.59-7.53
7.53-8.47
e
Oithonida
Sampling effort (log)
0-0.715
0.715-1.43
1.43-2.15
2.15-2.86
2.86-3.58
3.58-4.29
4.29-5.01
5.01-5.72
5.72-6.44
f
Poecilostomatoida
Sampling effort (log)
0-0.756
0.756-1.51
1.51-2.27
2.27-3.02
3.02-3.78
3.78-4.53
4.53-5.29
5.29-6.04
6.04-6.8
g
Euphausiacea
Sampling effort (log)
0-0.714
0.714-1.43
1.43-2.14
2.14-2.85
2.85-3.57
3.57-4.28
4.28-4.99
4.99-5.71
5.71-6.42
90°N
60°N
30°N
Eq
30°S
60°S
90°S
GM
60°E
120°E
180°
120°W
60°W
GM

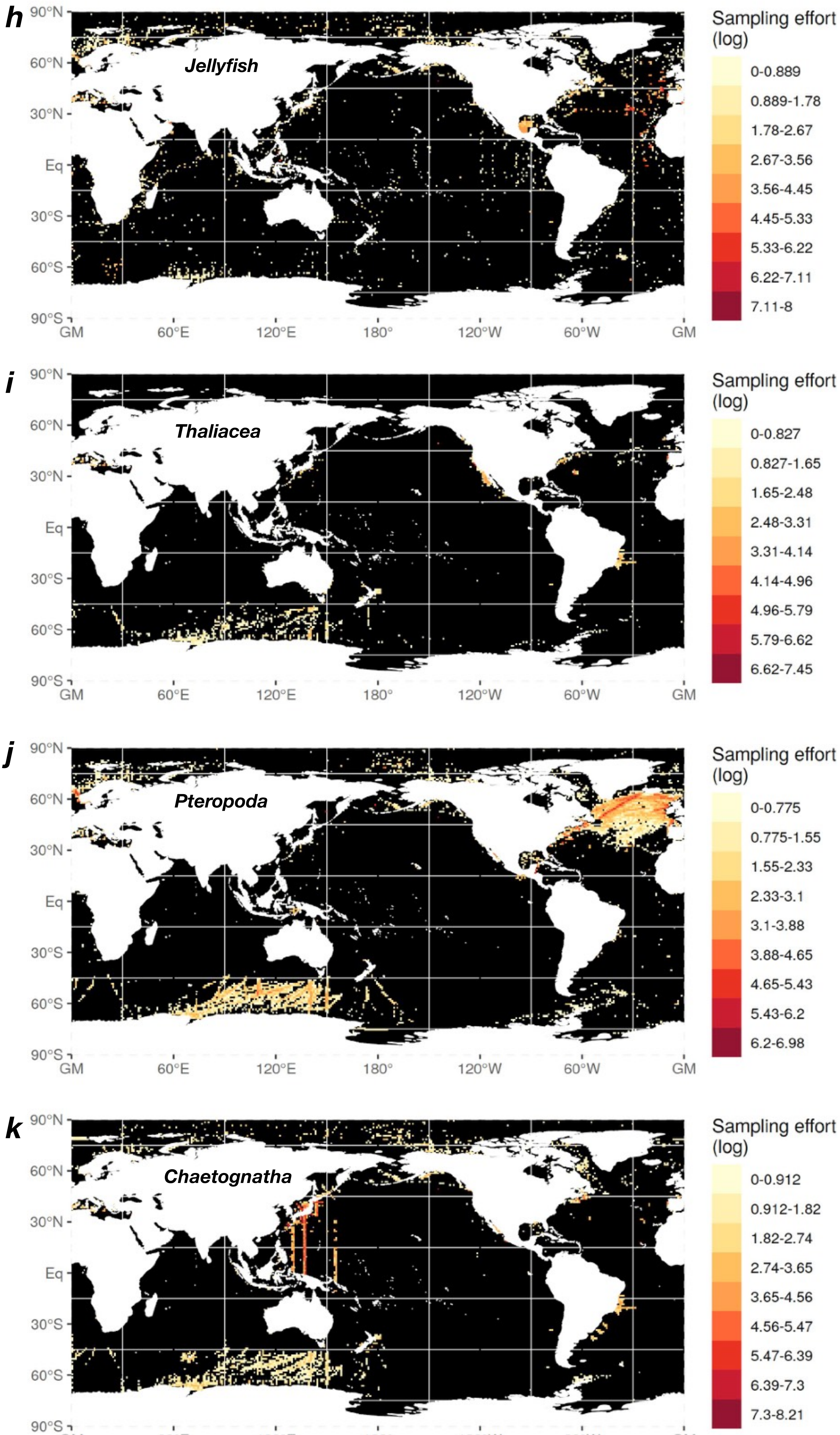
h
Jellyfish
90°N
60°N
30°N
Eq
30°S
60°S
90°S
GM
60°E
120°E
180°
120°W
60°W
GM
Sampling effort (log)
0-0.889
0.889-1.78
1.78-2.67
2.67-3.56
3.56-4.45
4.45-5.33
5.33-6.22
6.22-7.11
7.11-8
i
Thaliacea
90°N
60°N
30°N
Eq
30°S
60°S
90°S
GM
60°E
120°E
180°
120°W
60°W
GM
Sampling effort (log)
0-0.827
0.827-1.65
1.65-2.48
2.48-3.31
3.31-4.14
4.14-4.96
4.96-5.79
5.79-6.62
6.62-7.45
j
Pteropoda
90°N
60°N
30°N
Eq
30°S
60°S
90°S
GM
60°E
120°E
180°
120°W
60°W
GM
Sampling effort (log)
0-0.775
0.775-1.55
1.55-2.33
2.33-3.1
3.1-3.88
3.88-4.65
4.65-5.43
5.43-6.2
6.2-6.98
k
Chaetognatha
90°N
60°N
30°N
Eq
30°S
60°S
90°S
GM
60°E
120°E
180°
120°W
60°W
GM
Sampling effort (log)
0-0.912
0.912-1.82
1.82-2.74
2.74-3.65
3.65-4.56
4.56-5.47
5.47-6.39
6.39-7.3
7.3-8.21

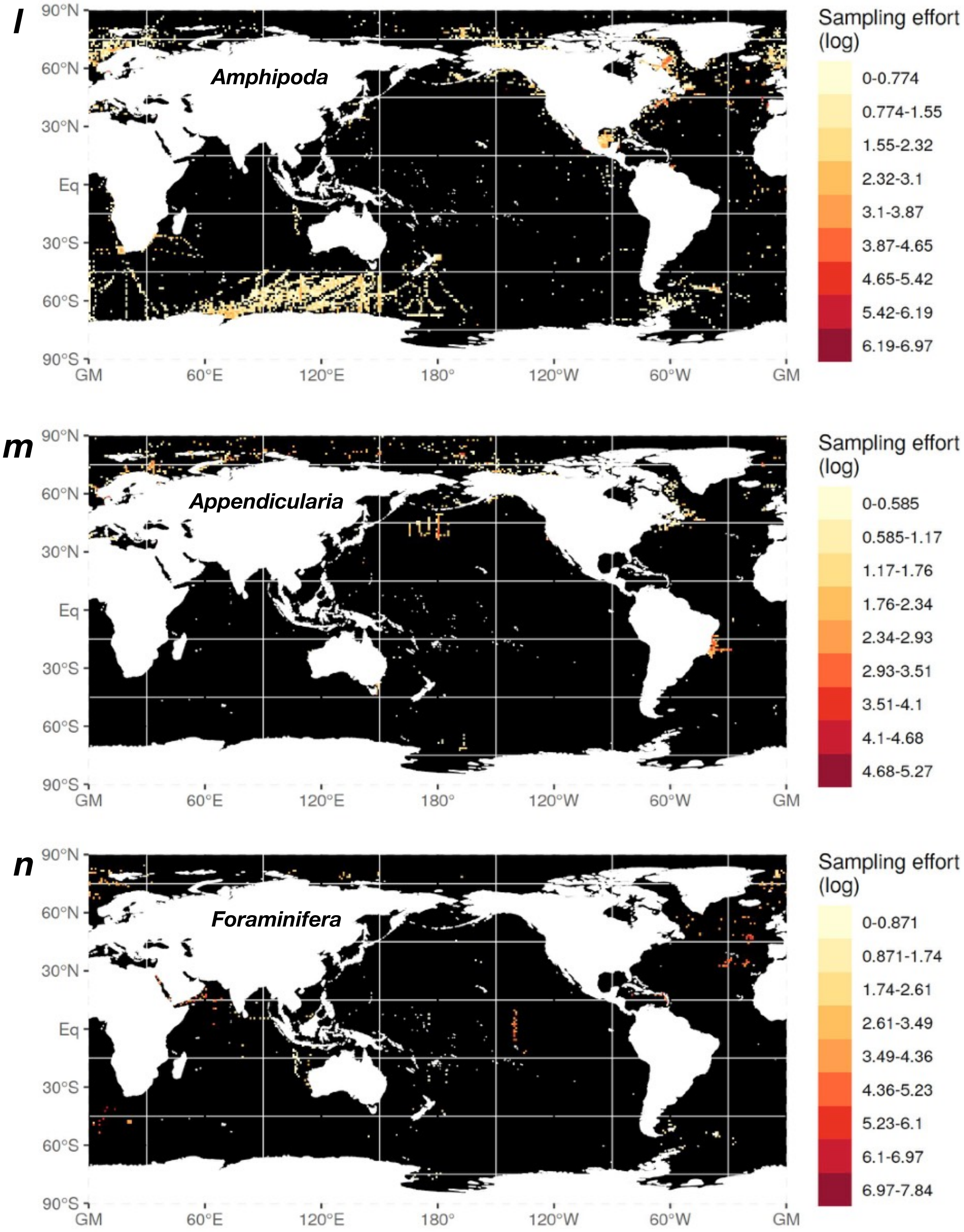
l
Amphipoda
90°N
60°N
30°N
Eq
30°S
60°S
90°S
GM
60°E
120°E
180°
120°W
60°W
GM
Sampling effort (log)
0-0.774
0.774-1.55
1.55-2.32
2.32-3.1
3.1-3.87
3.87-4.65
4.65-5.42
5.42-6.19
6.19-6.97
m
Appendicularia
90°N
60°N
30°N
Eq
30°S
60°S
90°S
GM
60°E
120°E
180°
120°W
60°W
GM
Sampling effort (log)
0-0.585
0.585-1.17
1.17-1.76
1.76-2.34
2.34-2.93
2.93-3.51
3.51-4.1
4.1-4.68
4.68-5.27
n
Foraminifera
90°N
60°N
30°N
Eq
30°S
60°S
90°S
GM
60°E
120°E
180°
120°W
60°W
GM
Sampling effort (log)
0-0.871
0.871-1.74
1.74-2.61
2.61-3.49
3.49-4.36
4.36-5.23
5.23-6.1
6.1-6.97
6.97-7.84
