## Supplementary figures and images for "Global gradients in species richness of marine plankton functional groups"

### Supplemental Figure 3

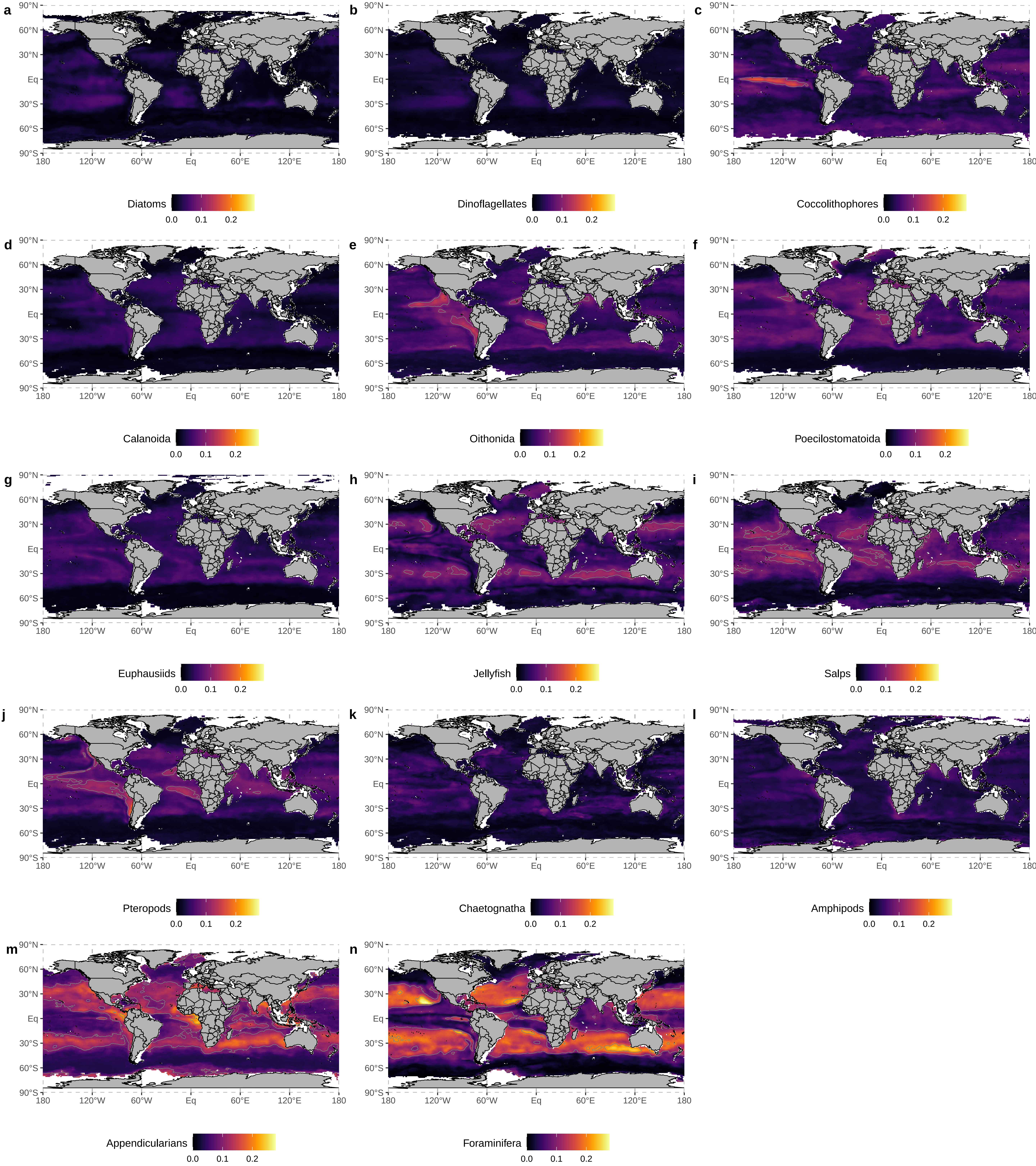
