## Supplemental Figure 4 for "Global gradients in species richness of marine plankton functional groups"

**Figure S4 : First four components of the principal component analysis (PCA) performed on the mean annual normalized species richness (SR) estimates of each plankton functional groups (PFG). The mean annual values of the environmental covariates and the proxies of megafauna diversity, plankton size structure and ecosystem functioning were added as supplementary variables projected on the PCA space a posteriori.**

**Figure S4.1**: Ordination plot showing how the mean annual normalized SR estimates of each PFG (black arrows) scored the first two principal components (PC1 and PC2) of the PCA. The length of the arrows indicate the strength of the variables' loadings on the principal components. The angles between the arrows indicate the level and direction of collinearity between the variables (90° angles reflect orthogonal variables meaning that are not collinear; 180° angles reflect a negative collinearity). Colored arrows indicate those variables that were projected a posteriori in the PCA space to examine their covariance with the mean annual SR estimates. Variables represented by blue arrows indicate those environmental covariates that were included in the species distribution models of the PFGs (but see Table S3). Variables represented by red arrows indicate those covariates that were never included in the species distribution models of the PFGs. The relative percentage of variance explained by each principal component is indicated in %.

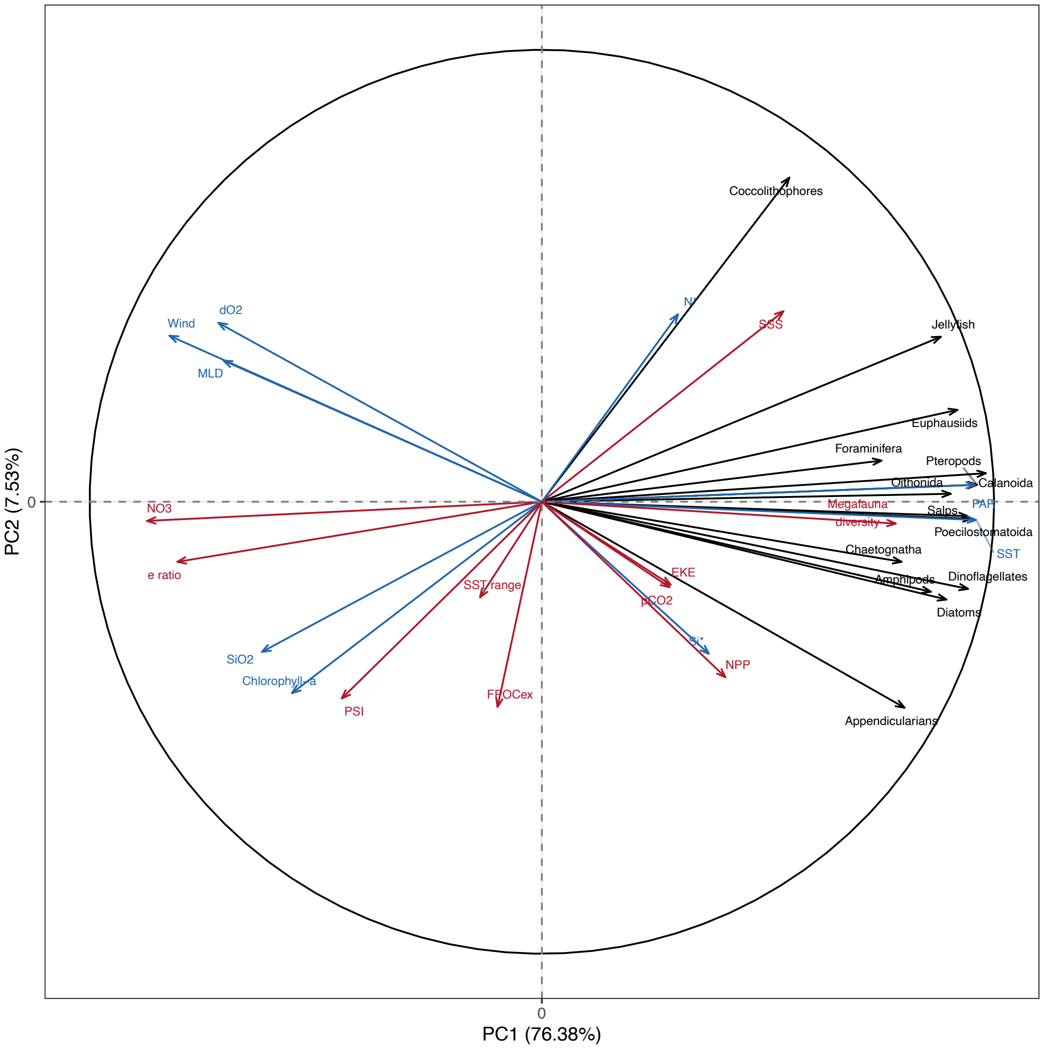

**Figure S4.2**: Same as Fig. S4.1, but with PC3 and PC4 of the PCA.

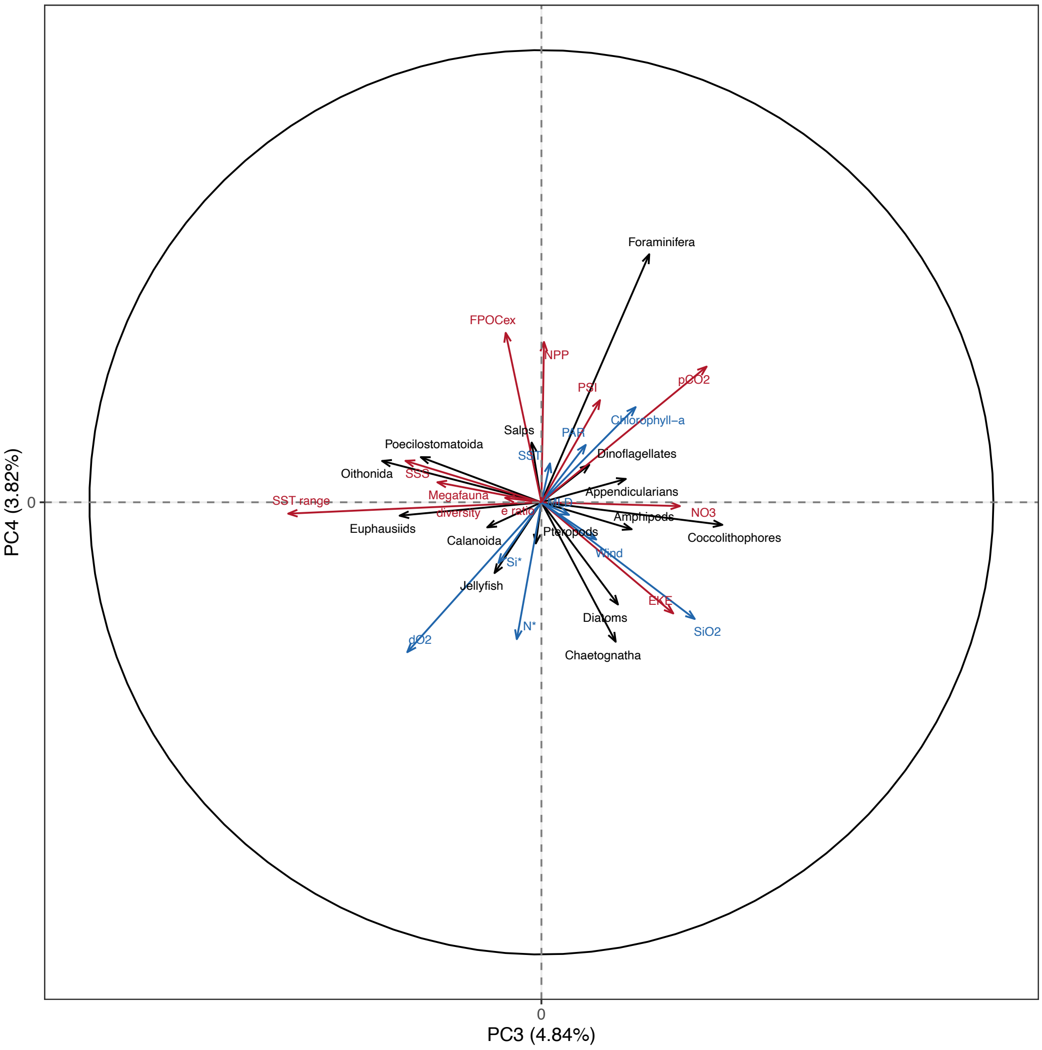
